# Size regulation of multiple organelles competing for a shared subunit pool

**DOI:** 10.1101/2020.01.11.902783

**Authors:** Deb Sankar Banerjee, Shiladitya Banerjee

## Abstract

How cells regulate the size of intracellular structures and organelles, despite continuous turnover in their component parts, is a longstanding question. Recent experiments suggest that size control of many intracellular assemblies is achieved through the depletion of a limiting subunit pool in the cytoplasm. While the limiting pool model ensures organelle size scaling with cell size, it does not provide a mechanism for robust size control of multiple co-existing structures. Here we propose a kinetic theory for size regulation of multiple structures that are assembled from a shared pool of subunits. We demonstrate that a negative feedback between the growth rate and the size of individual structures underlies size regulation of a wide variety of intracellular assemblies, from cytoskeletal filaments to three-dimensional organelles such as centrosomes and the nucleus. We identify the feedback motifs for size control in these structures, based on known molecular interactions, and quantitatively compare our theory with available experimental data. Furthermore, we show that a positive feedback between structure size and growth rate can lead to bistable size distributions arising from autocatalytic growth. In the limit of high subunit concentration, autocatalytic growth of multiple structures leads to stochastic selection of a single structure, elucidating a mechanism for polarity establishment.

## I. INTRODUCTION

Eukaryotic cells are composed of a wide diversity of macromolecular assemblies, from linear protofilaments to networks of cytoskeletal polymers and complex three-dimensional organelles such as the centrosomes and the nucleus. The cytoplasmic pool of proteins constitutes the building blocks for intracellular organelles, whose sizes are often commensurate with cell size. Despite continuous turnover in their component parts, intracellular organelles are maintained at a precise size through dynamic balance between subunit assembly and disassembly [1]. An outstanding challenge is to identify the design principles through which cells achieve robust size regulation of multiple co-existing structures that are assembled from a limiting pool of molecular building blocks in the cytoplasm.

Studies in recent years have focused on understanding the mechanisms for size control of individual cellular structures such as the eukaryotic flagella [2, 3], actin cables [4], mitotic spindles [5], centrosomes [6, 7], as well as the nuceloli [8] and the nucleus [9]. A simple model that explains size control of these dynamic structures is the ‘limiting pool’ model [1], where structures grow by depleting the pool of available subunits in the cytoplasm. As a result, growth rate of structures decreases with increasing assembly size, and a steady-state size is reached when the rate of assembly balances the rate of disassembly of incorporated material. Since organelle size is determined by the amount of available subunits in the cytoplasm, which in turn scales with cell size, the limiting pool model naturally captures the scaling of organelle size with cell size. However, the limiting pool fails to capture size regulation of multiple competing structures [10, 11], due to the absence of an underlying mechanism for sensing individual structure size. Failure of the limiting pool mechanism in determining the size of multiple structures suggests additional feedback design principles for organelle growth control. In this work we uncover the feedback motifs necessary for maintaining the size of multiple organelles, including the eukaryotic flagella, centrosomes, nuclei, as well as describe the mechanisms for co-existence multiple cytoskeletal filaments that assemble from a shared pool of monomers in the cellular cytoplasm [12, 13].

We begin by considering a deterministic description for the growth of *M* structures that incorporate material from *N*_av_ available subunits in the cytoplasm. Dynamics of size of the *i*^*th*^ structure (*i* = 1..*M*), *n*_*i*_ (expressed in the number of subunits), is given by:

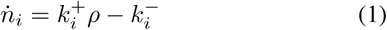

where *ρ* = *N*_av_*/V* is the concentration of subunits in the cytoplasm, *V* is the cell volume, and 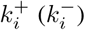 is the rate of assembly (disassembly) of the *i*^*th*^ structure. If the cytoplasmic concentration of subunits is maintained at a constant homeostatic value 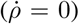 [14, 15], size control is not achieved except when *ρ* is fine tuned to a critical value *ρ* = *ρ*_*c*_ = *k*^−^*/k*^+^. Growth is unbounded for *ρ* > *ρ*_*c*_ and the assembly degrades for *ρ* < *ρ*_*c*_. By contrast, in the limiting pool mechanism, the total amount of subunits 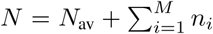 is constant. In this case, the assembled structure reaches a well-defined size *n* = *N* − (*k*^−^*/k*^+^)*V*, only for *M* = 1.

However, when multiple structures are assembled from a shared subunit pool (*M* > 1), (1) yields a system of under-determined rate equations with no unique solution for the steady-state size of the individual assemblies (see *SI: Section 1* for details). This indeterminacy manifests as large size fluctuations (Fig. S1) in a stochastic description [11], where building blocks can transfer between individual structures with no free energy cost. The underlying reason is that the limiting pool model does not provide a mechanism to sense the individual size of the structures. Rather, the limiting pool mechanism operates by sensing the size of the available subunit pool in the cytoplasm.

## MODEL FOR SIZE REGULATION OF MULTIPLE STRUCTURES GROWN FROM A SHARED SUBUNIT POOL

In the limiting pool model for *M* > 1, departure from size control of individual structures is manifested either as large anticorrelated size fluctuations (for identical growth rates), or the faster growing structure ends up incorporating all the subunits [11]. We therefore hypothesize that a negative feedback between size and the growth rate of individual structures might ensure robust size control. This motif can be realized when the net growth rate of individual structures decreases with increasing size. In fact, size-dependent assembly and disassembly rates have been reported in many cases of filament growth, including in Chlamydomonous flagella [3], microtubules [16, 17], as well as filamentous actin (F-actin) [18].

To elucidate the emergence of robust size control via size-dependent negative feedback, we consider the case of two assemblies growing from a shared pool of *N* subunits (Fig. 1A). At time *t*, the size of the *i*^*th*^ assembly (*i* = 1, 2) is given by *n*_*i*_(*t*), the number of incorporated subunits. We propose a minimal phenomenological model where the assembly rate of the *i*^*th*^ structure is 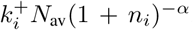, and the rate of disassembly is given by 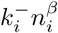. Here, 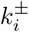 is the bare assembly(+) and disassembly(-) rates for the *i*^*th*^ structure, and *N*_av_ = *N* − (*n*_1_ + *n*_2_) is the total amount of available subunits. The coefficients *α* and *β* represent the strength of the autoregulatory feedback that can arise due to active molecular processes or the geometry of the structures grown. In this model, negative autoregulation of growth is ensured for *α* > 0 (assembly rate decreases with size) and/or for *β* > 0 (disassembly rate increases with size). We assume the subunit pool is well mixed in the cytoplasm such that subunit diffusion is much faster compared to the growth process. The stochastic system described by the assembly and disassembly processes yields the following steady-state probability distribution (see *SI: Section 2* for details),

**FIG. 1.**
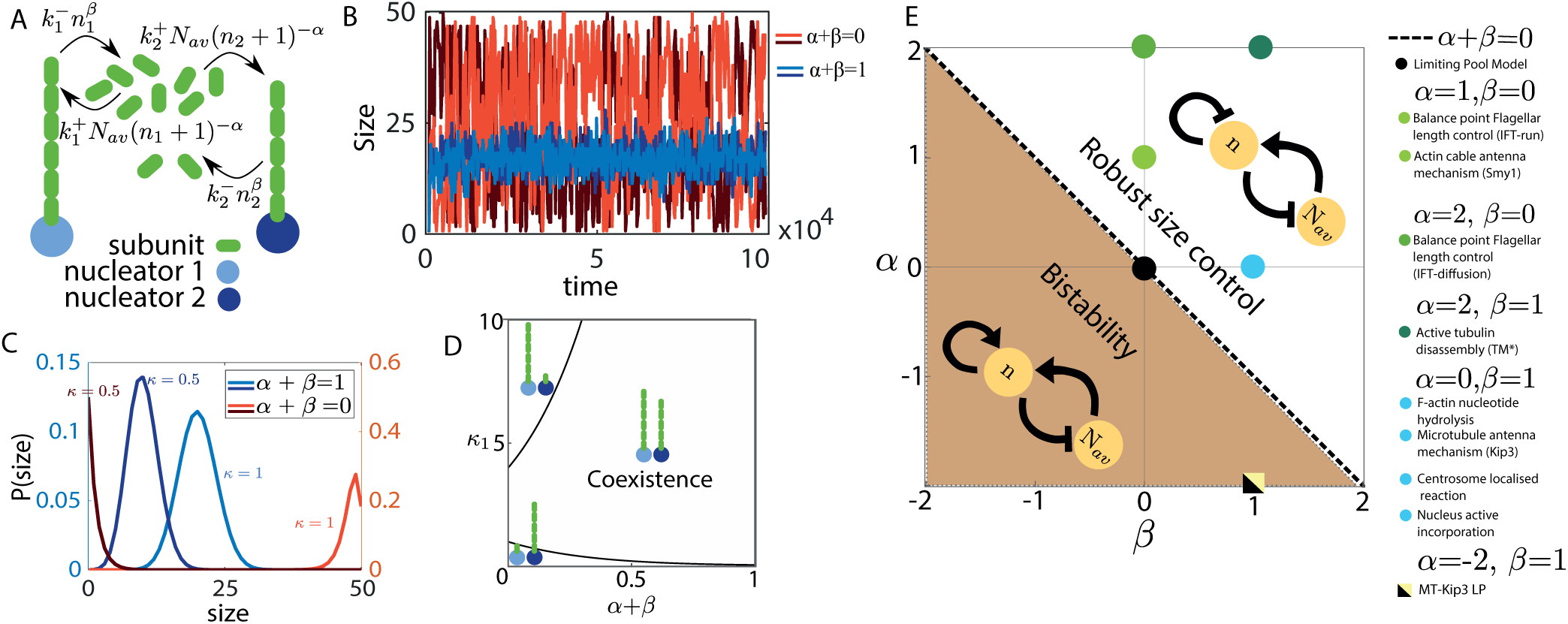
Size regulation of multiple structures grown from a shared subunit pool. (a) Schematic of two filaments growing from a shared pool of monomers where the assembly and disassembly rates depend on their individual size. (b) Time series for the size of two identical filaments grown from a shared pool of subunits in two limits of the model: *α* + *β* = 1 (light and dark blue lines), and *α* + *β* = 0 (light and dark red lines). (c) Size distribution of two competing structures with unequal growth rates (*κ*_1_ = 2*κ*_2_ = 1), for *α* + *β* = 1 (blue) and *α* + *β* = 0 (red). (d) Phase diagram showing co-existence of two competing structures over a broad range of parameter space, with *κ*_2_ = 2. For all results in (a-d) *V* = 1 and *N* = 50. (e) Phase diagram of the general growth model in *α*-*β* plane, showing the different regimes of size control. *α* + *β* > 0 defines the regime of negative autoregulation of growth which guarantees robust size control. Positive autoregulation of growth (*α* + *β* < 0) gives rise to bistable size distribution. The phase boundary *α* + *β* = 0 corresponds to the limiting pool model with constant assembly/disassembly rates, where there is no size regulation of individual structures. We map our model to size control mechanisms for a variety of intracellular structures, from linear filaments to 3D organelles.

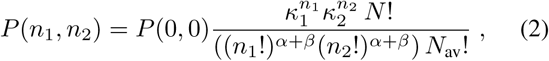

where 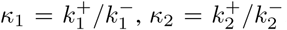, and *P* (0, 0) is a normalization constant.

The limit *α* + *β* = 0 corresponds to the limiting pool model with constant assembly and disassembly rates [1]. Lack of size regulation for *α* + *β* = 0 manifests as large anticorrelated size fluctuations of the two competing but identical structures (Fig. 1B). By contrast, for *α* + *β* > 0, there is an overall negative autoregulation of growth that ensures robust size control of multiple structures. To illustrate this, we numerically simulated (see Methods) the growth dynamics of two filaments competing for the same subunit pool, for the specific case *α* + *β* = 1 (Fig. 1B). The individual structures assume a well defined mean size (Fig. 1B), with the standard deviation in size fluctuations smaller compared to the mean (Fig. 1C). Furthermore, a negative autoregulation of growth rate ensures coexistence of multiple competing structures over a broad range of the parameter space (Fig. 1D).

To provide specific biological examples, the case (*α, β*) = (0, 1) exactly maps to length regulation of Microtubules and F-actin (see *SI: Section 3* and Fig. S2 for details), where the filament disassembly rate increases with increasing filament length. In the antenna mechanism for microtubule length control, the kinesin Kip3 associates with microtubule monomers, walks towards the plus end of the filament and detaches from the end by removing microtubule monomers [21]. Over time, Kip3 molecules accumulate near the plus-end, leading to an effective length-dependence of disassembly rate [22]. In the case of F-actin, chemical changes in subunit states, via nucleotide hydrolysis of bound monomers [23], can lead to length-dependent disassembly [18]. In addition, length-dependent F-actin disassembly could also arise through the action of the severing protein ADF/cofilin [24]. In all these cases, length-dependent disassembly via active molecular processes can stabilise the length of multiple filaments competing for the same monomer pool.

In the opposite scenario when *α* + *β* < 0, there is an overall positive feedback on growth (Fig. 1E), resulting in bistable size distributions. Bistable length regulation has been reported for microtubule-Kip3 systems [25], and we return to this interesting case later. In the sections that follow, we show that by tuning the model parameters *α* and *β*, we can quantitatively capture size control of diverse intracellular organelles, from the one-dimensional eukaryotic flagella to three-dimensional organelles such as centrosomes and the nucleus (Fig. 1E).

## LENGTH REGULATION OF MULTIPLE FLAGELLA ASSEMBLED FROM A SHARED POOL OF BUILDING BLOCKS

Flagellar growth in the biflagellate *Chlamydomonas reinhardtii* is a classic example of size regulation of multiple organelles assembled from a common cytoplasmic pool of building blocks [19, 20, 26]. Molecular mechanisms for flagellar length control remain an active area of research, with mathematical models suggesting that flagellum length dynamics is controlled by a length-dependent assembly process [2, 3, 27], or by a length-dependent disassembly mechanism [28].

*C. reinhardtii* flagella grow from a shared pool of tubulins, which are carried and assembled via intraflagellar transport (IFT) particles at the tip of the flagellum [3] (Fig. 2A). As the total amount of IFT particles on the flagellum remain constant over time [3], IFT density at the flagellar tip is a decreasing function of length. This leads to a length-dependent assembly rate for the flagellum, inversely proportional to the flagellum length [2], corresponding to the limit (*α, β*) = (1, 0) in our model (*SI: Section 4.A*, Fig. 1E). In addition, a length-dependent disassembly mechanism [28] would correspond to the limit (*α, β*) = (1, 1). Both these models fall within the general motif for size control via negative autoregulation of growth (Fig. 1E) that would guarantee robust length control for multiple flagella.

**FIG. 2.**
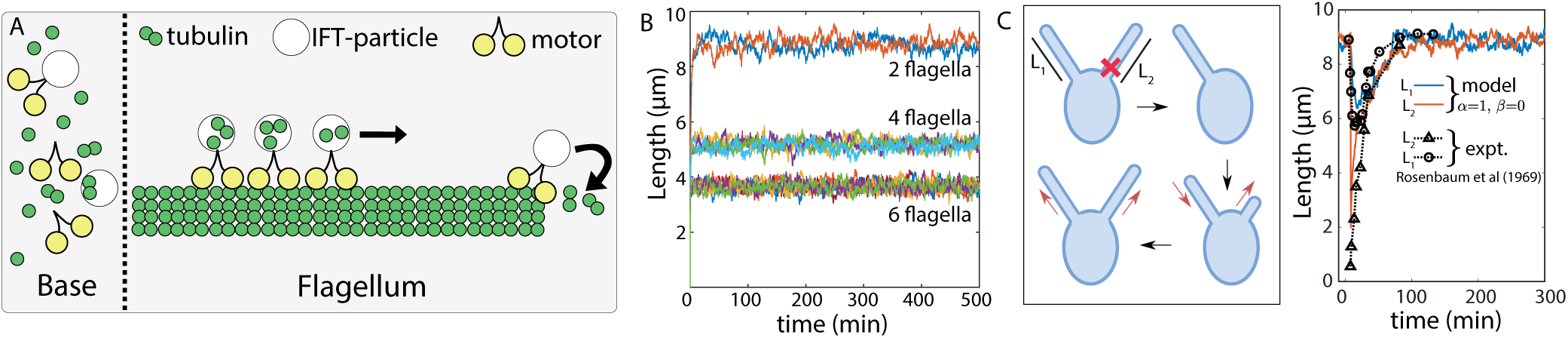
Length regulation of multiple flagella in *C. reinhardtii*. (a) Flagellum assembly in *C. reinhardtii* is regulated by IFT particles that incorporate tubulin dimers at the flagellum tip. This assembly process combined with the conservation of IFT amount in flagellum gives rise to an assembly rate that decreases with flagellum length. This mechanism corresponds to (*α, β*) = (1, 0) in our model. (b) Decrease in flagellar size with increasing number of flagella in mutant cells has been experimentally reported [10, 19]. Using stochastic simulations of our model, we show that increasing flagellum number for a fixed building block pool, results in flagellar size reduction at steady-state. For parameter values see *SI: Table S1*. (c) We model the flagellar re-growth experiment [20] where one of the two flagella is amputated and regrowth is observed. The intact flagellum starts shrinking immediately after the amputation, indicating a shared pool of building blocks. Our model with length-dependent assembly, (*α, β*) = (1, 0), quantitatively fits the experimental data for the length dynamics of the two flagella. For parameter values see *SI: Table S2*.

We use the balance-point model suggested by Marshall et al. [2, 3] to show that it is sufficient to regulate flagellar length in the multi-flagellate system in *C. reinhardtii*. A deterministic description of the system takes the form (*i* = 1, 2),

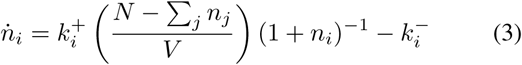

where *n*_*i*_ is the length of the flagellum in tubulin numbers, *V* is the cell volume, *N* is total tubulin amount in the cell. We use Gillespie algorithm (Methods) to simulate the stochastic system of multiple flagella grown from a shared pool of tubulins. Our simulations (Fig. 2B) capture the experimentally reported phenomena that mean flagellar size decreases with increasing the number of flagella in a cell [10, 19, 29]. We further use our stochastic model to simulate the flagellar regrowth experiment (Fig. 2C-left), where one of the two flagella is cut and re-growth is observed due to the production of new building blocks. We model this experiment (see *SI: Section 4.B* for details) by removing building blocks from one of the flagella at *t* = 0, resulting in rapid shrinkage in length of both the flagella. The timescale for fast shrinkage dynamics is governed by the rate constants *k*^±^. The slower process of length recovery is governed by the production rate of new subunits, which we calibrated by fitting our model to experimental data [19]. Our fitted model quantitatively captures the experimentally measured flagellar length dynamics (Fig. 2C-right).

## CENTROSOME SIZE CONTROL VIA LOCALISED ASSEMBLY AND DISTRIBUTED DISASSEMBLY

Having described how size-dependent negative feedback on growth can stabilise the length of multiple filamentous structures, we now turn to describing the mechanisms of size maintenance in three-dimensional organelles. Specifically we show that our proposed feedback motif for growth control (Fig. 1E) can describe centrosome size regulation during mitosis.

Centrosomes are membraneless spherical organelles consisting of a pair of centrioles at the center (Fig. 3A), surrounded by a porous scaffold-like structure [30] called the pericentriolic matter (PCM). In cells preparing to enter mitosis, the two centrosomes are spatially separated and grow by recruiting PCM material around the centrioles [6, 31–34]. While the mechanics of PCM assembly is a subject of ongoing debate [35], the molecular components for PCM growth must ensure robust size control of centrosomes during mitosis. Otherwise, small stochastic variations in the sizes of maturing centrosomes could amplify through the process of maturation, leading to errors in division ratio (Fig. 3A). Furthermore, experimental data show that centrosome size scales with cell size, through multiple rounds of cell divisions in the early *C. elegans* embryo, suggesting that centrosome size is determined by a limiting pool of building blocks [1, 6]. Since the limiting pool model cannot maintain the size of two structures competing for the same subunit pool (Fig. 1), additional feedback controls must be necessary for centrosome size regulation.

**FIG. 3.**
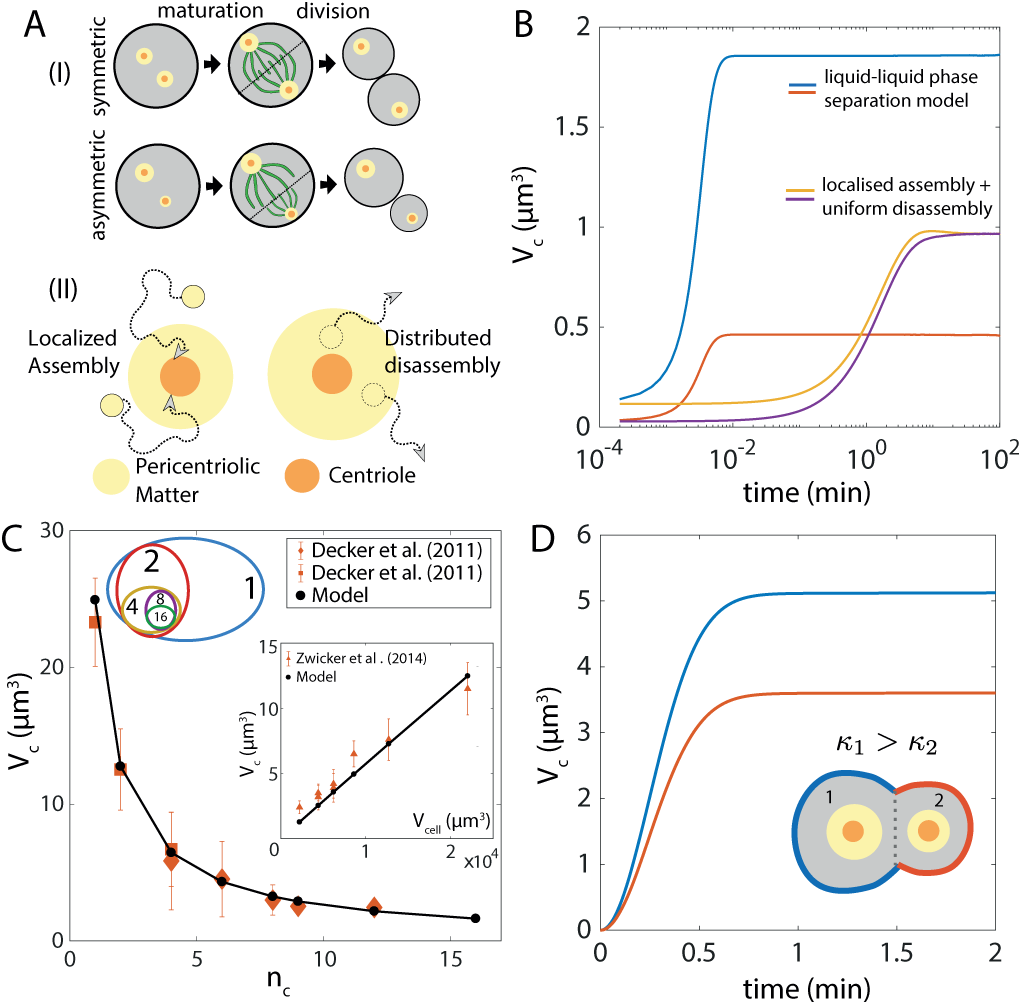
Size regulation of centrosomes. (a-I) Schematic of centrosome maturation in mitotic cells followed by cell division. Prior to maturation, spontaneous asymmetry in centrosome sizes can arise from stochastic variations. In the absence of size control, size asymmetry can amplify, resulting in asymmetric cell division. (a-II) Localised assembly around the centriole and disassembly throughout the pericentriolic matter generates a size-dependent disassembly rate which ensures robust size control. (b) Size dynamics of a pair of identical centrosomes that are initialised with unequal sizes. If assembly and disassembly occur uniformly throughout the volume, centrosome sizes will diverge (blue and red curves). By contrast, localised assembly and distributed disassembly can correct initial size errors to restore equal sized centrosomes (yellow and purple curves). For parameter values see *SI: Table S3,S4*. (c) Centrosome volume decreases with increasing centrosome number, *n*_*c*_, during *C. elegans* embryonic development, where the embryo rapidly divides into many cells with decreasing cell size. Model: solid black line, Experimental data: orange. Inset: Centrosome volume, *V*_*c*_, scales linearly with cell volume, *V*_cell_. For parameter values see *SI: Table S4*. (d) Since PCM assembly is controlled by the centriole, a centriole with higher nucleation rate will assemble PCM faster, leading to asymmetries in mitotic centrosome size. For parameter values see *SI: Table S5*.

Based on prior experimental observations [7, 34, 36, 37], we propose a kinetic model for the growth of spherical centrosomes that assemble PCM building blocks from a limiting cytoplasmic pool. PCM assembly is localized around the centriole, and disassembly can occur throughout the volume of the porous PCM matrix (Fig. 3A) [34, 36]. This design principle for organelle growth results in disassembly rate increasing with centrosome size, providing a size-dependent negative feedback control of growth. A deterministic description for this growth process is given by,

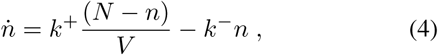

where *n* is the number of PCM building blocks incorporated in the centrosome, *k*^±^ are the bare assembly and disassembly rates, *N* is the total amount of the PCM building blocks, and *V* is the volume of the cytoplasmic pool. This growth mechanism directly maps to our general model for size-dependent growth with (*α, β*) = (0, 1) (Fig. 1E). Since *α* + *β* > 0, the model in (4) ensures robust size control of two centrosomes assembled from a shared resource pool.

We use the stochastic description of (4) to demonstrate size control of a pair of identical centrosomes (Fig. 3B). If there is an initial size difference between the two centrosomes, the model with localised assembly and distributed disassembly can correct the size error to restore equality of the size of centrosomes. By contrast, if assembly and disassembly occur uniformly throughout the volume, size asymmetries will amplify over time (Fig. 3B). Previous study [7] on centrosome growth via liquid-liquid phase segregation has emphasized the necessity of high centriole activity to form centrosomes. This model of centrosome as an active droplet with high centriole activity can be mathematically mapped to our kinetic model of localised assembly and distributed disassembly (see *SI: Section 5.A* and Fig. S3 for details). For small centriole activity, the liquid-liquid phase segregation model fails to control centrosome size (*SI: Section 5.A* and Fig. 3B). Aside from ensuring size control, our model for centrosome assembly can quantitatively capture the scaling of centrosome size with centrosome number (Fig. 3C) and cell volume (Fig. 3C-inset), as measured during early *C. elegans* embryonic development [6].

An early sigmoidal nature in centrosome growth has been identified as an essential feature of the regulatory growth mechanism [6]. This can naturally emerge from the way the different subunits are sequentially recruited in the PCM [34]. To this end, we consider a simple two-component growth model, where one component forms the scaffold structure and the other gets recruited to the scaffold. Thus the assembly rate of the second component depends on the scaffold size. This coupled assembly dynamics provides a positive feedback in the growth mechanism (see *SI: Section 5.B* for details). This results in an early sigmoidal growth of centrosome volume (Fig. 3D), as observed in experiments [6]. Our model predicts a positive correlation of the centrosome size with the centriole nucleation activity (∝ *k*^+^), as a higher nucleation rate will drive faster PCM assembly. This prediction can be related to the known dependence of centrosome size on centriole enzymatic activity [37, 38]. As a result, a centriole with higher *k*^+^ will assemble a larger centrosome (Fig. 3D), facilitating asymmetric cell division [39].

## NUCLEUS SIZE CONTROL BY SURFACE AND BULK GROWTH

Nucleus is a highly complex organelle, composed of two key components - the inner nucleoplasm (NP), surrounded by the outer nuclear envelope (NE). During nucleus growth, both the NP and NE components grow from their respective pool of building blocks (see *SI: Section 6* and Fig. S4). However, which one of these two components controls nucleus size and growth remains an open question [40]. Though there are many studies reporting the scaling of nucleus size with cell size [41–43], there are few to none theoretical models for nucleus size control. To this end, we present a simple two-component model for nucleus - an outer spherical shell representing NE, and an interior solid sphere representing NP. We demonstrate how the geometric design of NE and NP assembly lead to nuclear size regulation, and compare our model predictions with available experimental data.

We first consider a model for nucleus growth by nuclear envelope (NE) assembly, taking inspiration from a recent *in vitro* study in *Xenopus levis* egg extract [44], where nucleus growth is coupled to the growth of microtubule asters surrounding the nucleus [44–46]. Here, NE assembly occurs through active incorporation of nuclear membrane vesicles/fragments (building blocks of NE) by dynein motors moving along astral microtubule tracks (Fig. 4A). The microtubule aster is a collection of many dynamic microtubules surrounding the nucleus, where each individual filament grows from a cytoplasmic pool of tubulins (building blocks of microtubules) (Fig. 4A). The rate of NE assembly is proportional the size of the micro-tubule aster, as the number of available NE building blocks scales with the volume spanned by the the aster. As the NE grows in size maintaining a constant thickness, we assume that the NP volume expands accordingly to accommodate the increase in nuclear surface area (Fig. S4). The deterministic rate equations for the growth of one nucleus is given by

**FIG. 4.**
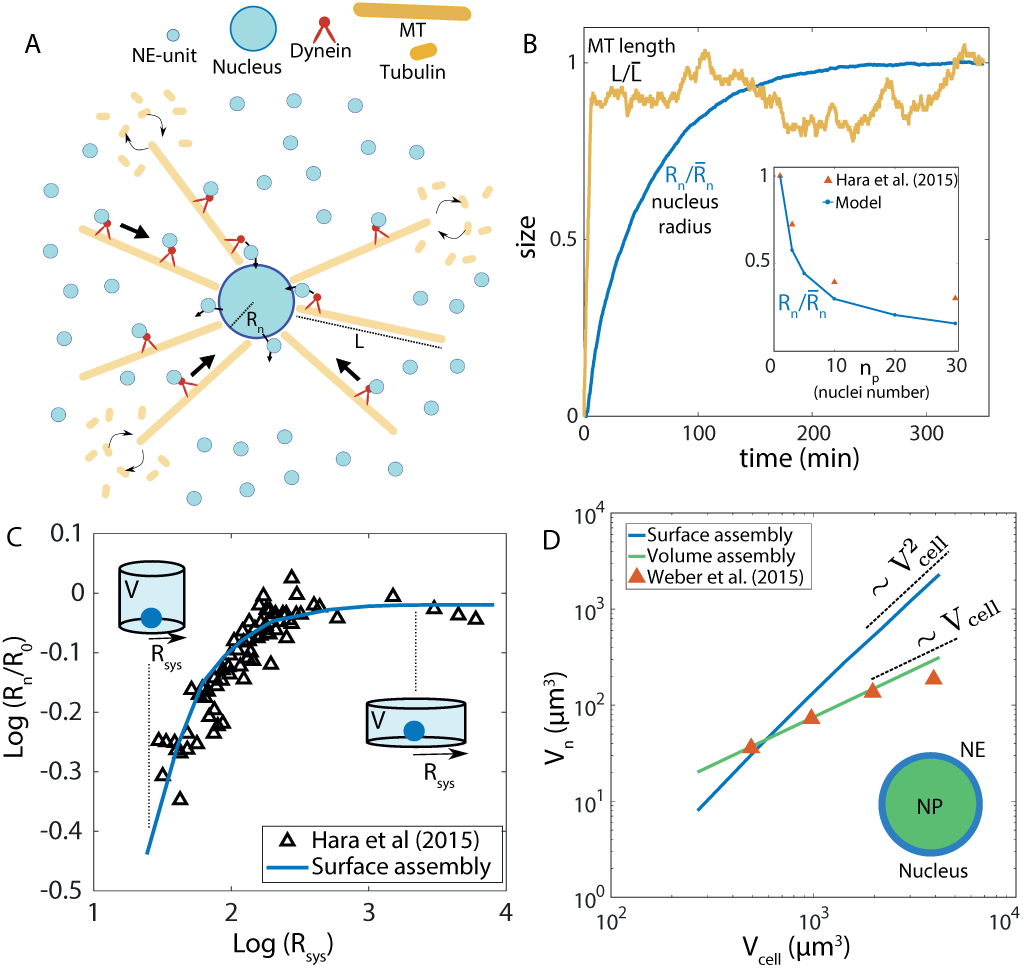
Nucleus size control. (a) Modelling the growth of nucleus, coupled to the dynamics of the astral microtubule structure. Building blocks for nuclear envelope (NE) are actively transported by dynein motors along the astral microtubules surrounding the nucleus. Filaments in the aster grow by incorporating tubulins from the cytoplasmic pool. (b) Dynamics of nucleus size (normalised radius *R*_*n*_) and single microtubule length (normalised). (Inset) Nucleus size decreases with increasing nuclei number in a given volume, in agreement with *in vitro* data [44]. (c) Effect of the size of confinement, *R*_sys_, on nucleus size *R*_*n*_, where *R*_*n*_ increases with increasing *R*_sys_, eventually saturating for large *R*_sys_. Solid line is model fit, and black triangles represent experimental data [44]. The confinement radius was increased while keeping the confinement volume constant, as in experiments [44]. (d) Scaling of nucleus volume, *V*_*n*_, with cell volume *V*_cell_. Theory predicts that size scaling is quadratic if *V*_*n*_ is controlled by the growth of NE, but the scaling is linear if *V*_*n*_ is regulated by NP assembly. The linear scaling fits quantitatively with the nucleus-to-cell size scaling measured in an eukaryotic cell [8]. For parameter values of models of nucleus growth by surface assembly (NE), volume assembly (NP) and microtubule growth see *SI: Table S6,S7,S8* respectively.

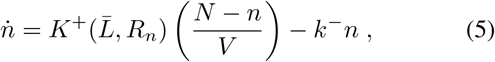

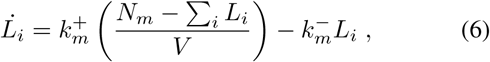

where *R*_*n*_ is the nucleus radius, *n* and *L*_*i*_ are the sizes of NE (in building block units) and the *i*^*th*^ microtubule filament, respectively. The total amount of tubulin and NE building blocks are given by *N*_*m*_ and *N*, respectively, and *V* is the cell (or system) volume. Here *k*^±^ and 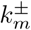 are the bare rates of assembly and disassembly for NE and microtubules, respectively.

Since assembly occurs at the surface, size of the nucleus is determined by the relation 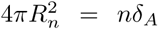, as *n* building blocks, each of area *δ*_*A*_, make up the NE. The size-dependent assembly rate is given by 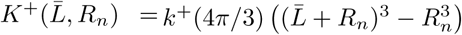, proportional to the volume accessible to the aster structure, and 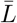 is the average length of the microtubules. Initially, the assembly rate induces a positive feedback on NE growth as 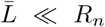. As assembly progresses, the filaments become longer to yield 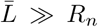, such that *K*^+^ becomes independent of *R*_*n*_. Disassembly occurs uniformly throughout the NE surface, yielding a size-dependent disassembly rate, *k*^−^*n*, which provides a negative feedback on NE growth. The later stages of NE growth can thus be mapped to our general growth model with (*α, β*) = (0, 1) (Fig. 1E), ensuring robust size control. Length dynamics of microtubules is implemented using the antenna model [21, 22], where the filament disassembly rate increases linearly with filament length (*SI: Section 3.A*).

We simulated the stochastic dynamics of nucleus growth using the model in (5) and (6), with the model parameters calibrated from *in vitro* data [44]. Both the filaments and the nuclear envelope reach a well-defined steady-state mean size (Fig. 4B). Initially, NE size increases rapidly due to size-dependent assembly rate (*K*^+^), whereas growth slows down at later times when *K*^+^ balances the rate of NE disassembly (Fig. 4B). Our model quantitatively captures the experimentally observed scaling between nucleus size and nuclei number, when multiple nuclei are assembled from a limiting pool of subunits (Fig. 4B, inset). The nuclear size, *R*_*n*_, decreases with increasing nuclei number, as expected from a limiting pool of building blocks. This coupled growth model of NE and microtubules can also explain the nucleus size dependence on the size of confinement as reported in *in-vitro* experiments [44]. Here, the microtubule growth is hindered by the confinement wall, and hence a smaller system size will generate a smaller aster structure. As a result, the assembly rate will be smaller giving rise to a smaller nuclear size (Fig. 4C). However, the maximum nucleus size is set by the steady-state length 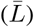 of the microtubule filaments (when all other conditions remain unchanged). Therefore, increasing the confinement size larger than 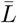 does not generate a larger nucleus (Fig. 4C). This can explain why an isolated nucleus grows to be larger in size than a group of closely packed nuclei in multinucleated cells. This is because the presence of neighbouring nuclei hinder microtubule growth [46], resulting in reduced nuclear size.

During nuclear growth in *Xenopus levis* egg extract, many NP proteins such as lamin-A, importin-*α* [9] are transported inside the nucleus and contribute to NP growth [47, 48]. To model growth of nucleus by NP assembly, we use the rate equations in (5) and (6), with the important difference that the size of nucleus is determined by the relation: 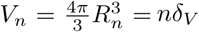, where *V*_*n*_ is the nucleus volume and *δ*_*V*_ is volume of individual NP subunits. Thus, when NP regulates nucleus size, *V*_*n*_ is proportional to the number of NP subunits incorporated,*n*. This model for nucleus growth by NP assembly does not alter the effect of confinement, but predicts the scaling relation, 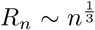. This scaling is different when nucleus size is regulated by NE assembly, where, 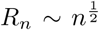 (Fig. 4D). This difference can be understood by relating NE-growth and NP-growth with growth of a spherical shell of constant thickness and the growth of a solid sphere, respectively. By comparing our simulation results with experimental data [8, 42], we find that the experimentally observed linear scaling of nuclear size with cell size cannot be achieved when nucleus size is purely regulated by NE assembly (Fig. 4D), but NP-growth model leads to *V*_*n*_ ∝ *V*_cell_. While our simplified description can capture experimentally reported nuclear size scaling and the effect of confinement on nuclear growth, the question of whether NP or NE set nucleus size cannot be answered without further experimentation. Our model predictions provide a possible way to test the underlying growth mechanism.

## BISTABLE SIZE REGULATION FROM AUTOCATALYTIC GROWTH

So far we discussed various examples of robust size control of multiple organelles via size-dependent negative feedback on growth (i.e., *α* + *β* > 0). In the opposite case of size-dependent positive feedback, *α* + *β* < 0, the dynamics are qualitatively different. For a single structure, positive feedback on growth implies an assembly (disassembly) rate that increases (decreases) with increasing structure size. This results in autocatalytic growth, where the structure initially grows at a faster rate due to the positive feedback, while eventually slowing down as the pool of building blocks is exhausted (Fig. 5A, inset).

**FIG. 5.**
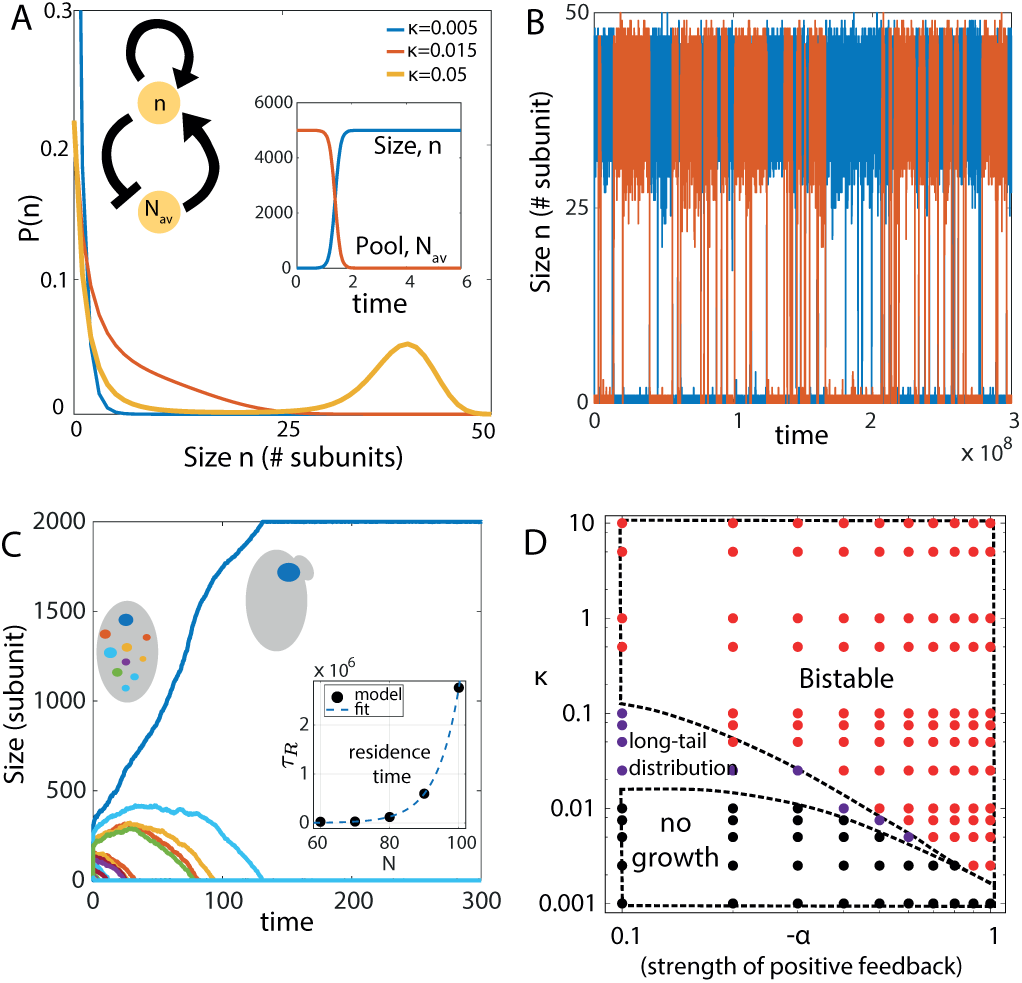
Bistable size distribution emerges from autocatalytic growth. (a) Probability distribution for the size of two identical structures assembled via autocatalytic growth. The structures do not grow at very low growth rates (*κ* = *k*^+^*/k*^−^), leading to an exponential distribution peaked at 0. At high *κ*, the size distribution is bistable. Inset: Single structure growth with size-dependent positive feedback exhibits sigmoidal growth, with the size eventually saturating when the subunit pool is completely exhausted. Parameters: *α* = −0.2, *β* = 0. (b) Dynamics of the size of the two structures in the bistable regime. Parameters: *κ* = 0.0025, *α* = −1.0, *β* = 0. (c) Stochastic selection of a single structure when multiple structures compete for a limiting subunit pool, in the limit *α* + *β* > 0 and high subunit concentration. Here, the residence time of a single structure becomes seemingly infinite, so the stochastically chosen large structure can remain stable for long timescales. Inset: Residence time (*τ*_*R*_) of a single structure increases exponentially with increasing concentration of subunits. Dashed line – exponential fit, solid circles – simulation data. Parameters: *N* = 2000, *κ* = 0.0125, *α* = −1, *β* = 0, and number of structures=20. (d) State diagram of the system, showing different growth regimes as a function of *κ* and the strength of positive feedback, −*α* (with *β* = 0). Change in *κ* induces a similar effect on the size dynamics as changing the overall concentration of subunits. By increasing *κ* at any non-zero value of −*α*, the system transitions from a “no-growth” state (black dots), to a “shoulder” state (purple dots) at intermediate *κ*, and finally a bistable state (red dots) for high *κ*. At very high *κ, τ*_*R*_ can be very large to effectively give rise to a single-structure. Increasing the strength of positive feedback promotes a bistable state at smaller *κ*. Parameters: *β* = 0, *V* = 1 and *N* = 50.

The result becomes less intuitive when there are two identical structures (with equal assembly/disassembly rates) competing for a shared subunit pool. When the pool consists of very few building blocks, the structures do not grow (Fig. 5A, blue). With sufficient building blocks, the structures initially grow at equal rates, but due to transient size differences arising from stochastic fluctuations, the bigger structure starts assembling faster than the smaller structure and ends up incorporating most of the building blocks. However, at this stage, stochastic fluctuations can make the larger structure lose enough building blocks to make a sudden transition to a smaller structure, while the other structure grows to be larger (Fig. 5B). Thus, at an intermediate concentration of building blocks, we get a bistable size distribution for two structures competing for a shared pool (Fig. 5A, yellow). There is an intermediate regime where the structures show large size fluctuations without bistable dynamics, and the size distribution exhibits a “shoulder” and a longer tail (Fig. 5A, red). Bistable size distribution has been reported in microtubule/kinesin-8 *in-vitro* systems [25], where kinesin-8 motors bind to the microtubules, walk towards the plus end and disassociate from the filament by removing a tubulin dimer [16, 21, 22]. This active disassembly depends on the motor concentration profile, which in turn depends on the filament length [17]. This can lead to a reduced concentration of motors at the tip of a longer filament, generating an effective positive feedback (with *α* = −2 and *β* = 1) and bistable length distribution (see *SI: Section 7* for details).

In the limit of high subunit concentration, a stochastically selected larger structure will consume all the building blocks to increase in size, but stochastic fluctuations may take a very long time to make the structures switch states. Depending on the strength of the positive feedback, growth rates and building block concentration, residence time of the bigger structure can be so large that the transition to a smaller structure may not occur within a realistic timescale. Thus an autocatalytic growth process (*α* + *β* < 0) can be used to direct the formation of a single structure. This phenomenon can be related to the process of polarity establishment in budding yeast, where the budding mechanism requires the formation of one single concentrated patch of the polarity protein Cdc42 that marks the budding location to initiate the subsequent process of budding [49]. Many studies have linked positive feedback in the process of Cdc42 patch formation via various physical mechanisms [50, 51] but the mechanism of polarity formation is not fully understood. Our model for autocatalytic growth of structures from size-dependent positive feedback is in good agreement with the previously stated mechanism of polarity establishment. In our case if we start with many structures only a few can survive to gain considerable size, but finally the stochastically chosen largest structure will win to form a single large structure while the other structures eventually die out (Fig. 5C).

Interestingly, Cdc42 proteins also exhibit size oscillations during polarity establishment in budding and fission yeasts [52, 53]. Our study shows that at an intermediate concentration of building blocks, multiple growing structures enter a bistable state (Fig. 5D and Fig.S5), where the residence time is small enough to promote transitions between large and small sized structure within experimentally relevant timescales. This is in good qualitative agreement with a recent study of Cdc42 oscillations where a decrease in overall protein amount promotes oscillations, generating many transient structures instead of one single large structure [53]. The negative feedback needed for these oscillations stems from subunit exchange with the limiting subunit pool.

## DISCUSSION

In this study, we uncovered the design principles for stable size regulation of intracellular structures and organelles in the noisy environment of the cell, where stochastic fluctuations may be significant. Our study reveals that a negative autoregulation of growth rate (*α* + *β* > 0) underlies robust size control of individual organelles and structures, when multiple of them compete for the same subunit pool. We demonstrate that our proposed feedback motif for size control is utilised by diverse subcellular structures, from one-dimensional filaments to three-dimensional organelles, by connecting our kinetic theory with known molecular processes in the cell. We show that our growth control model can also be utilised to assemble non-identical stable structures that may be important for cellular processes involving anisotropy and asymmetry. It is important to contrast our model with the limiting pool model for organelle growth control [1]. The latter provides a mechanism for organelle size scaling with cell size by sensing the subunit pool size, but fails to maintain the individual size of multiple competing organelles. The limiting pool model, however, succeeds in regulating the size of single structures because sensing the pool size is complementary to sensing the individual structure size.

It is natural to ask how cells regulate the size of intracellular structures when the subunit pool is not limited, but is maintained a constant homeostatic concentration. Our general growth model can also ensure size regulation of multiple structures in the case of subunit homeostasis when *α* + *β* > 0 (see *SI: Section 8* and Fig. S6 for details). Interestingly, subunit homeostasis does not lead to organelle size scaling with cell size and number of organelles (Fig. S6H), emphasizing the need for a limiting subunit pool to preserve organelle-to-cell size scaling. Subunit homeostasis can be important when structures are required to be maintained at a specific size regardless of cell size. In our proposed size-dependent growth model, it is possible to modulate the structure size to scale with cell size or be independent of cell size, by combining the features of subunit homeostasis and limiting pool model. Individual structure size would scale with the cell size when the cell volume *V* is smaller than 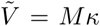 (Fig. S7 A-D), and saturates at 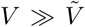 (Fig. S7 E-H), where *M* is the number of structures assembled. This result explains the experimentally observed independence of organelle size at larger cell sizes [8, 54] (see *SI: Section 9* and Fig. S7).

Subunit amount can increase during cell growth, as the abundance of many regulatory proteins increases with increasing cell size. It is therefore relevant to investigate the effect of cell growth (increasing cell size and subunit pool size) on organelle size control. If cell growth rate is faster compared to the assembly rate of the structures, the limiting pool mechanism can maintain transiently stable sizes for multiple structures grown from a shared subunit pool (Fig. S8A). But when cell growth is much slower than structure growth, then the limiting pool model fails to regulate the size of individual structures, exhibiting large size fluctuations (Fig. S8B). In this case, a negative feedback autoregulation of growth rate is required to achieve robust size control (Fig. S8 C-D) (see *SI: Section 10*).

Our proposed model for organelle growth control is general enough to capture size regulation for diverse subcellular structures and organelles that employ distinct molecular mechanisms to achieve size-dependent negative feedback control of growth rates. We map various existing size control mechanisms to our model by changing only two parameters, including in three-dimensional organelles such as centrosomes and nucleus. Specifically, we show how a membrane-less organelle like centrosome can self-assemble in multiple numbers while maintaining stable sizes, where a passive liquid-liquid phase separation model for centrosome assembly fails to achieve robust size control. We also demonstrate how centrosomes can maintain stable size differences under competition, which could serve as a precursor to asymmetric cell division. In the case of nuclear size regulation, our model is able to capture the experimentally reported nuclear-to-cell size scaling, and the effect of confinement where nucleus grows smaller in size when the local nucleus density is higher. In both cases of centrosomes and nuclear growth control, we provide quantitative comparisons of our model with available experimental data.

In the presence of positive feedback between structure size and growth rate, we uncover novel phenomena such as bistable size distribution where structures dynamically fluctuate between a larger and a smaller assembly. Interestingly, the transition rate from the larger to the smaller structure becomes vanishingly small when the subunit pool is large, giving rise to a single stochastically chosen large structure that is maintained for very long timescales. This elucidates a mechanism of spontaneous symmetry-breaking and polarity establishment, which is relevant for understanding the mechanism of bud formation in *S. cerevisiae* from the autocatalytic growth of Cdc42 clusters.

## METHODS

### Stochastic growth simulations

We use the Gillespie algorithm [55] to simulate the stochastic growth of one or multiple structures from a common pool of subunits. At any time *t* the Gillespie algorithm uses two random variables drawn from an uniform distribution (*r*_1_, *r*_2_ ∈ 𝒰(0, 1)), and the instantaneous propensities for all of the possible reactions to update the system in time according to the defined growth law. The propensities of the relevant reactions, i.e., the assembly and disassembly rates of the *i*^*th*^ structure are given by 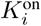 and 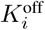 respectively. For our growth model these propensities are functions of subunit pool size (*N*) and structure size (*n*_*i*_),

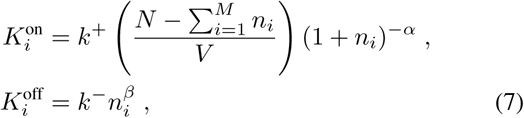

where we are considering growth of *M* structures from a shared pool. The Gillespie algorithm computes the time for the next reaction at *t* + *τ* given the current state of the system (i.e., the propensities for all reactions) at time *t* where *τ* is given by-

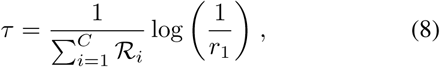

where ℛ_*i*_ is the propensity of *i*^*th*^ reaction and *C* is the total number of all possible reactions which is equal to 2*M* in our case. The second random variable *r*_2_ is used to select the particular reaction (*j*^*th*^ reaction) that will occur at *t* + *τ* time such that

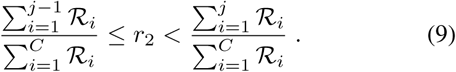

The condition for the first reaction (*j* = 1) is 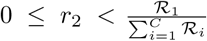 The two steps defined by Eq. 8 and Eq. 9 are used recursively to compute the growth dynamics in time.

## ACKNOWLEDGEMENTS

SB acknowledges support from the Royal Society (grant URF/R1/180187). SB and DB acknowledge support from the Human Frontiers Science Program (HFSP) grant RGY0073/2018. SB and DB thank Nikola Ojkic and Michael Murrell lab for critical comments on the manuscript. DB thanks Suman G. Das and Rituparno Mandal for useful discussions.

## Notes

#### Summary of Updates

Supplemental file uploaded

