## Supplementary Text for "Size regulation of multiple organelles competing for a shared subunit pool"

### I. LIMITING POOL MODEL

During the growth of one or multiple structures via limiting pool mechanism, cell maintains a constant total amount of subunits $N$ (no net production or degradation of subunits). As structures grow in size, the subunit pool depletes (so density of subunits decreases) and consequently the assembly rate decreases. So the growth dynamics for the $i^{th}$ structure when $M$ number of structures are grown from a shared subunit pool, can be written as

$$\dot{n_i} = k_i^+ \left( \frac{N - \sum_{i=1}^{M} n_i}{V} \right) - k_i^- \tag{1}$$

where $k_i^{\pm}$ is the assembly and disassembly rates for $i^{th}$ structure and $n_i$ is the size of the $i^{th}$ structure in number of monomers/subunits. The available amount of subunit at any given time is given by $N_{av} = N - \sum_{i=1}^{M} n_i$. Now for a single structure ($M = 1$) growth the equation becomes

$$\dot{n} = k^+ \left( \frac{N - n}{V} \right) - k^- \tag{2}$$

with the parameters having usual meaning as before. We can calculate the steady state (i.e., $\dot{n} = 0$) size from this to be

$$n^* = \left( \rho_0 - \frac{1}{\kappa} \right) V \tag{3}$$

where overall subunit density $\rho_0 = \frac{N}{V}$ and $\kappa = \frac{k^+}{k^-}$. This result from the deterministic description is identical to the mean size calculated from the respective stochastic description. If the overall concentration of subunits is maintained at a constant value (i.e., $\rho_0$ is independent of $N$ and $V$) we obtain a linear scaling of structure size with cell size (Fig. 1c).

Now if we consider multiple structures growing from a shared pool with equal growth rates ($k_i^{\pm} = k^{\pm}$ for all $i$), then we get a system of under-determined equations at steady-state (i.e., $\dot{n_i} = 0$ for all $i$) leads us to

$$k^+ \left( \frac{N - \sum_i n_i}{V} \right) - k^- = 0 \tag{4}$$

where the equations for $n_i$'s are identical and any combination of sizes that satisfies $\sum_i n_i = \left( \rho_0 - \kappa^{-1} \right) V$ is a solution for the above equation. This hints at the failure of size control for the growth of multiple structures from a limiting subunit pool.

Notably the total size of all structures is robustly controlled (Fig. 1d-e). A stochastic description for the growth of multiple structures captures the failure of size control as studied recently [1]. In the case of two structures, the size distributions are uniform. With many more structures, the size distribution converges to an exponential function, such that the standard deviation of size fluctuations is of the same order as the mean size (Fig. 1f).

### II. SIZE-DEPENDENT GROWTH MODEL

Here we discuss a model for size-dependent growth, where the growth mechanism senses the individual structure size via a negative autoregulatory feedback. We show that this size-dependent growth model can ensure robust size control of multiple structures. We first consider a deterministic description of the system, given by

$$\dot{n_i} = k_i^+ \left( \frac{N - \sum_i n_i}{V} \right) (1 + n_i)^{-\alpha} - k_i^- n_i^{\beta} \tag{5}$$

where $\alpha$ and $\beta$ quantify the strengths of size-dependent feedback, which might arise from geometry of the growth process or other active molecular processes that involve a promoter or a damper of growth. In the case of flagellar growth [2, 3] and microtubule growth [4] the intra-flagellar transport (IFT) particles and kinesin motors can be identified as such additional active elements. This deterministic description captures the mean sizes accurately for $(\alpha, \beta) = (1, 0)$ or $(\alpha, \beta = (0, 1)$, and the system of equations are not underdetermined for $M > 1$, unlike the limiting pool model.

The stochastic description for two structures of size $n_1$ and $n_2$, growing from a shared pool of total subunits $N$ within a system of unit volume ($V = 1$) is given by the following master equation for the joint probability-

$$\frac{\mathrm{d}P(n_1,n_2,t)}{\mathrm{dt}} = k_1^+(N-(n_1+n_2-1))n_1^{-\alpha}P(n_1-1,n_2,t) + k_2^+(N-(n_1+n_2-1))n_2^{-\alpha}P(n_1,n_2-1,t)$$
$$+k_1^-(n_1+1)^{\beta}P(n_1+1,n_2,t) + k_2^-(n_2+1)^{\beta}P(n_1,n_2+1,t)$$
$$-\left(k_1^+(N-(n_1+n_2))(n_1+1)^{-\alpha} + k_2^+(N-(n_1+n_2))(n_2+1)^{-\alpha} + k_1^-n_1^{\beta} + k_2^-n_2^{\beta}\right)P(n_1,n_2,t) \tag{6}$$

here $P(n_1,n_2,t)$ is the probability of finding the system with the structures in size $n_1$ and $n_2$ at time $t$. The steady state probability (i.e. when $\frac{dP}{dt}=0$) can be found by solving this equation using the following detailed balance conditions

$$k_1^-n_1^{\beta}P(n_1,n_2) = k_1^+(N-(n_1+n_2-1))n_1^{-\alpha}P(n_1-1,n_2)$$
$$k_2^-n_2^{\beta}P(n_1,n_2) = k_2^+(N-(n_1+n_2-1))n_2^{-\alpha}P(n_1,n_2-1)\ , \tag{7}$$

which yields,

$$P(n_1,n_2) = \left(\frac{\kappa_1(N-(n_1+n_2-1))}{n_1^{\alpha+\beta}}\right)P(n_1-1,n_2)$$
$$= \left(\frac{\kappa_1(N-(n_1+n_2-1))}{n_1^{\alpha+\beta}}\right)\left(\frac{\kappa_2(N-(n_1+n_2-2))}{n_2^{\alpha+\beta}}\right)P(n_1-1,n_2-1) \tag{8}$$

The above relation can be used iteratively to determine the steady-state joint probability distribution

$$P(n_1,n_2) = \left(\frac{\kappa_1^{n_1}\kappa_2^{n_2}}{(n_1!)^{\alpha+\beta}(n_2!)^{\alpha+\beta}}\right)\left(\frac{N!}{(N-(n_1+n_2))!}\right)P(0,0) \tag{9}$$

where $P(0,0)$ is the probability of finding both the structures at zero size and it can be calculated using the normalization condition. This exact solution for the steady-state joint distribution may or may not be obtained in a functional form, but we can easily compute the marginals $P(n_1)$ and $P(n_2)$ by summing over the other variable -

$$P(n_{1,2}) = \sum_{n_{1,2}=0}^{N}\left(\frac{\kappa_1^{n_1}\kappa_2^{n_2}}{(n_1!)^{\alpha+\beta}(n_2!)^{\alpha+\beta}}\right)\left(\frac{N!}{(N-(n_1+n_2))!}\right)P(0,0). \tag{10}$$

The above sum can be numerically computed to evaluate the discrete size distributions. In the limit $\alpha+\beta=0$, we recover the result for the joint distribution in the limiting pool model,

$$P(n_1,n_2) = \kappa_1^{n_1}\kappa_2^{n_2}\left(\frac{N!}{(N-(n_1+n_2))!}\right)P(0,0). \tag{11}$$

### III. FILAMENT LENGTH CONTROL BY LENGTH-DEPENDENT DISASSEMBLY

#### A. Antenna model for microtubule length control

Microtubule filaments grow via the addition of tubulin dimers as the subunits. In presence of Kip-3 motors, the disassembly rate of microtubule filaments becomes length dependent, as found experimentally [5, 6]. The motors bind to the filaments and move towards the minus end and detaches a tubulin dimer from the filament. The amount of motors walking on a filament increases with the increasing filament length thus the disassembly rate increases with filament length. This length dependence of disassembly rate was found to be linear [5]. Hence the disassembly rate for the $i^{th}$ microtubule of size $n_i$ can be written as $k^-n_i$ and the deterministic description of the growth for microtubule antenna model becomes

$$\dot{n_i} = k^+\left(\frac{N-\sum_{i=1}^{M}n_i}{V}\right) - k^-n_i \tag{12}$$

where $M$ is the number of microtubule filaments growing from a shared pool of tubulin. This description maps exactly to our model (Eq. 5) with $\alpha = 0,\ \beta = 1$. Here $k^+$ and $k^-$ are the bare assembly and disassembly rates, respectively, and $k^-$ can be an increasing function of motor density [5], which can tune the mean microtubule filament size

$$n^* = \frac{N}{\left(M + \frac{k^-}{k^+}V\right)}. \tag{13}$$

#### B. Nucleotide hydrolysis for F-actin length control

It has been reported that treadmilling-like growth of actin filaments can lead to an effective length-dependent disassembly rate, via nucleotide hydrolysis of the the bound monomers [7]. Here we take a simple case of treadmilling-like growth of one actin filament where the bound G-actin monomers can be in either of two states – ATP-bound state or ADP-bound state. We consider a minimal model where the filament length increases via binding of ATP-bound monomers at the barbed end with binding rate $k^+$, and length decreases only via unbinding of monomers from the pointed end. Tthe unbinding rate depends on the monomer state, and we denote $k_b^-$ and $k_u^-$ as the unbinding rates for ATP-bound and ADP-bound monomers respectively. If $P_b(x,t)$ and $P_u(x,t)$ define the probabilities of finding the monomer at position $x$ (in the frame of the filament with origin at barbed end) and time $t$ in the ATP-bound and the ADP-bound states, respectively, then we can express $P_b(x, t + \Delta t)$ as

$$P_b(x, t+\Delta t) = k^+\Delta t P_b(x-\Delta x, t) - k^+\Delta t P_b(x,t) + w_1 \Delta t P_u(x,t) - w_2 \Delta t P_b(x,t) \ , \tag{14}$$

which leads to the following master equation –

$$\partial_t P_b(x,t) = -k^+\,\partial_x P_b(x,t) + w_1 - (w_1 + w_2 + 1)P_b(x,t) \ , \tag{15}$$

where $w_1$ and $w_2$ are the transition rates of monomers going from ADP-bound state to the ATP-bound state and *vice versa*. Assuming the system reaches a steady-state ($\partial_t P_b = 0$) at filament length $\bar{L}$, we can solve for $P_b(x)$ as

$$P_b(x) = A + (1-A)e^{-\lambda x} \tag{16}$$

where $A = \frac{w_1}{w_1+w_2+1}$ and $\lambda = \frac{w_1+w_2+1}{k^+}$. Here we have employed the assumption that only ATP-bound monomers are binding to the barbed end which lead us to the following boundary condition: $P_b(0) = 1, P_u(0) = 0$. At steady-state, the effective disassembly rate ($k_d^{\text{eff}}$) at the pointed end and the assembly rate at the barbed end should be the same, such that

$$k^+ = k_d^{\text{eff}} = k_b^- P_b(\bar{L}) + k_u^-(1 - P_b(\bar{L})) \ . \tag{17}$$

Using Eq.16 we can then compute the length dependence of the disassembly rate

$$k_d^{\text{eff}}(\bar{L}) = k_u^- + (k_b^- - k_u^-)\left(A + (1-A)e^{-\lambda \bar{L}}\ .\right) \tag{18}$$

The steady-state filament length $\bar{L}$ is given by

$$\bar{L} = \lambda^{-1}\ln\left(\frac{1-A}{P-A}\right) , \tag{19}$$

where $P = \frac{k^+ - k_u^-}{k_b^- - k_u^-}$. In the limit of small $\lambda$, i.e. $w_1 + w_2 + 1 \ll k^+$, we get

$$k_d^{\text{eff}} = C_0 + C_1\bar{L} \tag{20}$$

where $C_0 = k_b^-$ and $C_1 = \lambda(k_u^- - k_b^-)(1-A)$. This limit approximately corresponds to our size-dependent growth model with $(\alpha, \beta) = (0, 1)$.

### IV. FLAGELLAR GROWTH

#### A. Length-dependent assembly

Flagellar growth in *Chlamydomonas reinhardtii* is controlled by the intra-flagellar transport (IFT) process. IFT-tubulin complex proteins are carried to the flagellum tip where they incorporate the tubulin subunit in the flagellum contributing to flagellar growth. The total amount of IFT-complex remains unchanged with flagellum length [3], hence the IFT current at the flagellum will decrease with increasing flagellum length. The elongation rate was reported to scale inversely with the flagellum length [3]. We can write the assembly rate as $k^+ N_{av}(1+n_i)^{-1}$, giving rise to the following deterministic description for flagellar growth –

$$\dot{n_i} = k^+ \left(\frac{N - \sum_{i=1}^{M} n_i}{V}\right)(1+n_i)^{-1} - k^- \ . \tag{21}$$

This model exactly maps to our size-dependent growth model with $\alpha = 1$, $\beta = 0$.

#### B. Flagellar regrowth model

We use the stochastic version of Eq. 21 to simulate the well known *long-zero* experiment for flagellar regrowth, where one of the two flagella is cut at $t = t_0$ and the length dynamics for both the flagella are measured. Upon severing of one of the flagella, the intact flagellum starts shrinking initially whereas the amputated flagellum starts growing in response to the cut [8]. Both the flagella start growing after some time, eventually reaches a steady state-length that is similar to the original length of the flagellum [8]. The regrowth of the damaged flagellum is limited by the production of new subunits as inhibition of protein production (using cycloheximide) produces two flagella with the same but smaller size [8].

In the simulation, we start with the normal growth of two flagella (with lengths $L_1$ and $L_2$), and let the flagella size to reach a steady-state following the stochastic dynamics for the description given in Eq. 21. We model the amputation by letting $L_2 \to 0$ which results in reduction in the total number of building blocks $N = N_{av} + L_1 + L_2$, such that $N \to N - L_2$. Subunits are produced at a rate $r_p$ to replenish the available subunit pool such that $N_{av}(t) = N_{av}(t_0) + \delta N(t)$. Here the production process is taken to be deterministic for simplicity and it is given by

$$\delta\dot{N} = r_p(\Delta_N - \delta N) \ , \tag{22}$$

where $\Delta_N (= L_2)$ is the lost amount of subunits during the flagellum cut and the initial condition is $\delta N(t_0) = 0$. We used a least-squared minimization process to determine the production rate by fitting our model with the experimental data of the *long-zero* experiment. The final results are shown using a rescaled time $t - t_0$. Our results quantitatively reproduce the dynamics of flagellar length in the long-zero experiment.

### V. CENTROSOME GROWTH MODEL

#### A. Self-assembly vs Liquid-liquid phase segregation model

The maturation process of mitotic centrosomes is a complex process involving multiple molecular players. The physical process of forming a large aggregate of pericentriolic matter (PCM) around the centriole during maturation is not well understood. Recent studies have suggested the role of liquid-liquid phase segregation in the centrosome maturation [9] but it still remains a highly debated issue [10]. Here we discuss centrosome growth using the model of liquid-liquid phase segregation adapted from Zwicker *et al* [9], to show how it can be mapped to our kinetic self-assembly model. We specifically discuss PCM droplet growth in the limit where phase segregation is strong and growth is limited by chemical reaction (i.e., fast diffusion of PCM components).

The growth of the centrosome (considered as a liquid droplet) volume $V$ is given by

$$\frac{dV}{dt} = (k\phi_1^A - k_{BA})V + Q\frac{\phi_1^A}{\psi_-} + k_{AB}\frac{\phi_0^A V_c}{N\psi_-} \tag{23}$$

where $\phi^A$ and $\phi^B$ are the volume fractions for the soluble ($A$) and the phase segregated ($B$) forms of the PCM components, respectively, and $\psi_-$ is the volume fraction of $B$ inside the droplet. Here $k_{AB}$, $k_{BA}$ and $k$ are the reactions rates for $A \to B$, $B \to A$ and $AB \to B$ respectively. The bulk volume fraction of the $A$ and $B$ forms, away from the droplet, is given by $\phi_0^A$ and $\phi_0^B$ respectively and $\phi_1^A$ is the volume fraction of $A$ inside the droplet. The chemical activity of the centriole, volume of the cell, and the number of centrioles growing are given by $Q$, $V_c$ and $N$ respectively.

We neglect spontaneous production of phase segregated form $B$ away from the centriole ($k_{AB} = 0$), and the volume fraction of $B$ inside the droplet is considered to be unchanged, i.e., $\psi_-$= constant. The volume fractions of $A$ in the bulk and inside the droplet are given by

$$\phi_0^A = \bar{\phi} - \psi_- \left(\frac{NV}{V_c}\right) \tag{24}$$

$$\phi_1^A = (1 - \psi_-)\phi_0^A \tag{25}$$

where $\bar{\phi}$ is the average volume fraction of the total PCM material. We define $\bar{\phi} = \frac{V_A+V_B}{V_c} = \frac{N_{\text{tot}}}{V_c}$, where $N_{\text{tot}}$ is the total amount of PCM material that remains constant during the droplet growth. We also write $\psi_- = \frac{n_B \delta_v}{V}$, where $n_B$ and $\delta_v$ are the number of $B$ subunits inside droplet and characteristic volume of each subunit respectively. Now we can rewrite $\phi_0^A = \frac{N_{av}\delta_v}{V_c}$ and $\phi_1^A = (1-\psi_-)\left(\frac{N_{av}\delta_v}{V_c}\right)$ where $N_{av} = N_{\text{tot}} - n_B$ is the available amount of subunits that can contribute to the droplet growth. Finally using the definition of $\psi_-$ we get the $n_B$ dynamics given by

$$\left(\frac{\delta_v}{\psi_-}\right)\frac{dn_B}{dt} = \frac{dV}{dt} \tag{26}$$

and this simplifies to

$$\frac{dn_B}{dt} = (1-\psi_-)\left(kn_B + \bar{Q}\right)\left(\frac{N_{av}\delta_v}{V_c}\right) - k_{BA}n_B \tag{27}$$

where $\bar{Q} = \frac{Q}{\delta_v}$. This above description then can be rewritten as

$$\dot{n}_B = (C_0 + C_1 n_B)\left(\frac{N_{av}}{V_c}\right) - C_2 n_B \tag{28}$$

where $C_0 = (1-\psi_-)Q$, $C_1 = (1-\psi_-)k\delta_v$ and $C_2 = k_{BA}$. In the case of $C_0 \ll C_1$ (passive phase separation limit) this growth description maps to our size dependent growth model with $\alpha = -1, \beta = 1$. Thus this growth mechanism will lie on the limiting pool line, $\alpha + \beta = 0$, which does not guarantee size control for multiple structures. We compare the growth of two centrosomes via passive liquid-liquid phase separation (LLPS) and by our localised assembly mechanism to demonstrate that in passive LLPS model the stochastic fluctuations in size will lead to different sizes of centrosomes, while our self-assembly model can ensure robust size control of yielding two centrosomes of same average size in steady state (Fig. 3). In the limit $C_0 \gg C_1$ where the phase segregation is controlled by centriole chemical activity ($Q$), the LLPS model results are very similar to our localized assembly model, and is able to provide robust size control for multiple centrosomes.

#### B. Two-component model

During mitotic centrosome maturation it has been reported [11] that some of the PCM proteins start assembling a scaffold-like structure around the centriole (component: $n_a$) and other components assemble on this scaffold (component: $n_b$). Thus the assembly of the second kind of components depends on the scaffold formation. For simplicity we take the assembly rate of $n_b$ to be a linearly increasing function of $n_a$ (i.e., faster assembly rate for larger scaffold). The deterministic description of the two component growth can be written as

$$\dot{n}_a = k_a^+\left(\frac{N^a - n_a}{V}\right) - k_a^- n_a \tag{29}$$

$$\dot{n}_b = k_b^+\left(\frac{N^b - n_b}{V}\right) n_a - k_b^- n_b \tag{30}$$

where $k_{a,b}^{\pm}$ is the assembly(+) and disassembly(-) rates for the $a$ and $b$ components. The volume of the centrosome is then given by $V_{\text{cen}} = (n_a + n_b)\delta V$. This two component coupled growth shows the early time sigmoid growth (emerging from the $n_a$ dependent positive feedback in $n_b$ growth) that has been observed in experiments during centrosome maturation. We note that this positive feedback does not play any major role in the robust control of centrosome size against stochasticity or noise, but perhaps it might be important for other physiological functions such as asymmetric cell division.

### VI. NUCLEUS GROWTH CONTROL

Nucleus is a membrane-bound organelle, whose structure, dynamics and constitutive molecular components are tightly regulated for proper cell functionality. Here we consider a simplified description of nucleus composed of two components- the outer nuclear envelope (NE) and the interior nucleoplasm (NP). We present a simple theory of nucleus growth where we investigate two hypotheses for size control: A. nucleus size is set by the NE surface area, and B. nuclear size is set by NP volume. We assume that when the size is set by the NE, NP volume expands accordingly to accommodate the increase in NE surface area, and *vise versa*.

#### A. Nuclear size control by nuclear envelope growth

Recent *in vitro* experiments [12] in *Xenopus* egg extract have reported the growth of nucleus from the assembly of nuclear envelope (NE). The size of the nuclear envelope grows from the assembly of the NE-subunits (Fig. 4a) which is carried by dyenin motors walking on microtubule tracks [12]. These microtubule tracks form an aster-like structure around the nucleus [12]. In the case of the nuclear size regulation by the NE size the radius of the nucleus is given by

$$R_n = \left(\frac{n\delta_A}{4\pi}\right)^{\frac{1}{2}} \tag{31}$$

where $n$ and $\delta_A$ are the number of NE-subunits incorporated into the nuclear envelope and the area of a typical NE-subunit, respectively. Addition of a NE-subunit increases the NE area by $\delta_A$ and changes the nuclear radius according to the above equation. Here we assume that the nucleoplasm volume increases with NE surface area increase to completely fill the volume enclosed by the nuclear envelope. Thus the dynamics of nuclear envelope size set the size of the nucleus. It is important to note that due to the above relationship between nuclear radius and NE-size, the nucleus volume scales as $V_n \sim n^{\frac{3}{2}}$.

#### B. Nuclear size control by nucleoplasm growth

The nucleus size depends on various nucleoplasmic proteins such as lamin-A, importin-$\alpha$ [13] and nucleoplasmin (Npm2) [14]. These proteins might act as the limiting components for nucleoplasm growth. In the case of nuclear size being set by nucleoplasm growth from NP-subunit assembly (Fig. 4b), the radius of the nucleus is given by

$$R_n = \left(\frac{3n\delta_V}{4\pi}\right)^{\frac{1}{3}} \tag{32}$$

where $\delta_V$ is the volume of a typical NP subunit and $n$ denotes the total number of such subunits incorporated in the nucleoplasm. Here we assume that the nuclear envelope grows accordingly to enclose the nucleoplasm. Thus the volume of the nucleoplasm sets the size of the nucleus. This leads to the scaling prediction $V_n \propto n$, which is different than the previous case of size control by NE-growth. In this model of nuclear growth by NP assembly, we considered a similar case of mictrotubule-assisted assembly of NP-subunits via active transport and NP disassembly rate to be proportional to the NP volume.

It is important to note that the difference in nucleus volume scaling in the above two models leads to differences in scaling of the nucleus volume with cell volume (as in steady state $n \sim V_{\text{cell}}$). Nucleus size scaling in cells (not in embryos where the mechanism can be different) can be better explained by a model of nuclear growth with NP assembly. While it is possible that both the mechanisms play vital roles in nucleus growth control, it is likely that the most scarce subunit components set the size of the nucleus. More experimental studies are needed to differentiate between these two proposed growth mechanisms.

### VII. BISTABLE SIZE DISTRIBUTION FROM AUTOCATALYTIC GROWTH: THE KIP-3 AND TUBULIN LIMITING POOL MODEL

In a recent study by Rank *et al* [15], it was demonstrated how Kip-3 motors that disassemble microtubules subunits (the tubulin dimers) from the plus-end give rise to bistable microtubule length distribution. Here, we show how this mechanism can be mapped to our model in the limit $\alpha + \beta < 0$, where we find bistable size distribution due to autocatalytic growth from size-dependent positive feedback and negative-feedback from the limiting pool. In Rank *et al*'s study, the assembly rate is given by

$$K_{\text{on}} = \gamma_0 \left(c_T - \frac{L}{\delta_L V}\right) \tag{33}$$

where $L$ is the filament, $V$ is the cell/system volume, $c_T = N/V$ is the initial subunit density, and $\delta_L$ is the subunit length. The constant $\gamma_0$ is the same as the bare assembly rate, $k^+$, in our description. Disassembly occurs at the microtubule plus-end, when a Kip-3 motor reaches the end and falls off with a subunit. Thus the disassembly rate is given by

$$K_{\text{off}} = \rho_+(t)\,\Delta \tag{34}$$

where $\rho_+(t)$ is the probability that the plus-end has a motor at time $t$, and $\Delta$ is the dissociation rate in the presence of a motor. The estimate of the probability leads to the following expression [15] for the disassembly rate

$$\rho_+(t)\,\Delta = \frac{w_A L}{\delta_L} - \left(\frac{(w_A + w_D) w_A L^2}{2\delta_L^2 v}\right)\ , \tag{35}$$

where $w_A$ and $w_D$ are the rates at which Kip-3 motors bind and unbind from the filament, respectively, and $v$ is the velocity of motors walking on the microtubule. The length dynamics is given by

$$\dot{L} = \delta_L \left( K_{\text{on}} - K_{\text{off}} \right). \tag{36}$$

The length here is $L = n\delta_L$ where $n$ is the number of tubulins in the filament. The subunit density is $c_T = \frac{N}{V}$, where $N$ is the total subunit pool size and $V$ is the system size. Now the length dynamics can be re-written in terms of the subunit number as follows,

$$\dot{n} = k^+ \left( \frac{N_{av}}{V} \right) + C_1 n^2 - C_2 n \ , \tag{37}$$

where $C_1 = \frac{w_A(w_A+w_B)}{2v}$ and $C_2 = w_A$. The above equation approximately maps to our size-dependent growth model with $\alpha = -2$ and $\beta = 1$, belonging to the regime $\alpha + \beta < 0$. We have shown that in this regime our model exhibits bistable size distribution, which agrees with the reported results of Rank et al. Interestingly this model points out a regime where the positive feedback is very weak (i.e., $\frac{k^+ N_{av}}{V} \gg C_1 n^2$), giving rise to autocatalytic growth and bistability. In this regime, microtubule growth can be mapped to our model with $\alpha = 0, \beta = 1$, leading to robust length control for multiple microtubule filaments competing for a limiting pool.

### VIII. CONSTANT SUBUNIT CONCENTRATION MODEL: FAILURE OF SIZE CONTROL AND ITS RECOVERY

When the cell maintains a constant subunit concentration, $\rho$, the growth dynamics for $M$ number of structures growing from a shared pool can be written as

$$\dot{n}_i = k_i^+ \rho - k_i^- \tag{38}$$

where $k_i^{\pm}$ is the assembly and disassembly rates for $i^{th}$ structure and $n_i$ is the size of the $i^{th}$ structure in number of subunits. The structures will keep growing without bound when $\dot{n}_i > 0$. This unbound growth occurs above a critical subunit density $\rho_c$ given by $\dot{n}_i = 0$ -i.e., $\rho_c = k_i^- / k_i^+$. Below this critical density the structures do not grow at all. Recovery from this failure in size control occurs when the growth process has size-dependent negative feedback

$$\dot{n}_i = k_i^+ \rho (1 + n_i)^{-\alpha} - k_i^- n_i^{\beta} \tag{39}$$

where $\alpha + \beta > 0$. We take a simple case of two structures (Fig. 6a) where the corresponding master equation for the growth of two structures of size $n_1$ and $n_2$ is given by

$$\begin{aligned} \frac{\mathrm{d}P(n_1, n_2, t)}{\mathrm{d}t} &= k_1^+ \rho n_1^{-\alpha} P(n_1 - 1, n_2, t) + k_2^+ \rho n_2^{-\alpha} P(n_1, n_2 - 1, t) \\ &+ k_1^- (n_1 + 1)^{\beta} P(n_1 + 1, n_2, t) + k_2^- (n_2 + 1)^{\beta} P(n_1, n_2 + 1, t) \\ &- \left( k_1^+ \rho (n_1 + 1)^{-\alpha} + k_2^+ \rho (n_2 + 1)^{-\alpha} + k_1^- n_1^{\beta} + k_2^- n_2^{\beta} \right) P(n_1, n_2, t) \ , \end{aligned}$$

where $P(n_1, n_2, t)$ is the probability that the two structures have sizes $n_1$ and $n_2$ at time $t$. The steady-state probability is obtained by solving the above master equation using the following detailed balance conditions –

$$\begin{aligned} k_1^- n_1^{\beta} P(n_1, n_2) &= k_1^+ \rho n_1^{-\alpha} P(n_1 - 1, n_2) \\ k_2^- n_2^{\beta} P(n_1, n_2) &= k_2^+ \rho n_2^{-\alpha} P(n_1, n_2 - 1) \ , \end{aligned} \tag{40}$$

which yields –

$$P(n_1, n_2) = \left( \frac{\kappa_1 \rho}{n_1^{\alpha+\beta}} \right) \left( \frac{\kappa_2 \rho}{n_2^{\alpha+\beta}} \right) P(n_1 - 1, n_2 - 1) \tag{41}$$

where $\kappa_1 = \frac{k_1^+}{k_1^-}$ and $\kappa_2 = \frac{k_2^+}{k_2^-}$. We can use this relation to iteratively compute the steady-state probability distribution:

$$P(n_1, n_2) = \left( \frac{\kappa_1^{n_1} \kappa_2^{n_2} \rho^{n_1+n_2}}{(n_1!)^{\alpha+\beta} (n_2!)^{\alpha+\beta}} \right) P(0, 0) \ , \tag{42}$$

where $P(0,0)$ is the probability of finding both the structures at zero size and it can be calculated using the normalization condition. This exact solution of steady state joint distribution of size may or maynot be written in a closed functional form, but we can easily calculate the marginals $P(n_1)$ and $P(n_2)$ by summing over the other variable –

$$P(n_{1,2}) = \sum_{n_{1,2}=0}^{\infty} \left( \frac{\kappa_1^{n_1} \kappa_2^{n_2} \rho^{n_1+n_2}}{(n_1!)^{\alpha+\beta} (n_2!)^{\alpha+\beta}} \right) P(0,0). \tag{43}$$

We compute this sum numerically to calculate the discrete size distributions. We can easily see that taking $\alpha = 0, \beta = 0$ we get a probability distribution that is not normalizable signifying unbounded growth and the non-existence of a steady-state size distribution (unless $\kappa_{1,2}$ or $\rho$ is very small to give rise to non-growing structures) (Fig. 6b-c). It is important to note the role of the nucleator in this model, which enables us to attribute a non zero transfer rate from the $P(0,0)$ state to $P(0,1)$ or $P(1,0)$. With $\alpha + \beta > 0$ we show that we recover from the unbounded growth, and obtain size control of multiple structures with or without competition (Fig. 6d-g). But this growth mechanism does not provide access to information on cell/system size. Thus it is not possible to get organelle size scaling with cell size or organelle number, when subunit concentration is maintained at a constant value (Fig. 6h).

### IX. ORGANELLE-TO-CELL SIZE SCALING

Size of intracellular organelles often scale with cell size for proper physiological functionality, but it may also be desirable for organelle sizes not to scale with cell size. Our proposed model for organelle growth control by size-dependent negative feedback, enables us to tune the organelle-to-cell size scaling. To illustrate this, we take the examples of $\alpha = 0,\ \beta = 1$ and $\alpha = 1,\ \beta = 0$. In the following, we assume that all structures grow with the same bare assembly and disassembly rates, i.e., $k_i^{\pm} = k^{\pm}$. This assumption leads to identical steady-state sizes ($n^*$) for all the $M$ number of growing structures, such that $\sum_i n_i = Mn^*$.

For $\alpha = 0,\ \beta = 1$ the deterministic rate equations are given by,

$$\dot{n}_i = k^+ \left( \frac{N - \sum_{i=1}^{M} n_i}{V} \right) - k^- n_i \ , \tag{44}$$

which leads to the steady-state solution,

$$n^* = \frac{\kappa N}{\kappa M + V} \ , \tag{45}$$

where $\kappa = k^+/k^-$, $V$ is the cell volume and $M$ is the number of structures. The overall subunit density, $\rho_0 = N/V$, is a constant independent of cell size $V$ and total pool size $N$. We can thus rewrite the steady-state size as,

$$n^* = \frac{\kappa \rho_0 V}{\kappa M + V}. \tag{46}$$

In the limit $\kappa M \gg V$, the organelle size scales with linearly with cell size as $n^* \sim \rho_0 V/M$ (Fig. 7c), and scales inversely with number of organelles assembled (Fig. 7d). By contrast, when $\kappa M \ll V$ then $n^* \sim \kappa \rho_0$, i.e., the organelle size is independent of cell size and M (Fig. 7a,b).

For $\alpha = 1,\ \beta = 0$ the deterministic rate equations are,

$$\dot{n}_i = k^+ \left( \frac{N - \sum_{i=1}^{M} n_i}{V} \right) (1 + n_i)^{-1} - k^- \ , \tag{47}$$

which leads to the steady-state solution

$$n^* = \frac{(\kappa \rho_0 - 1)V}{\kappa M + V}. \tag{48}$$

As before, in the limit $\kappa M \gg V$ we get linear scaling of organelle size with cell size, $n^* \sim \frac{(\kappa\rho_0 - 1)V}{\kappa M}$ (Fig. 7g), and inverse scaling with the number of organelles (Fig. 7h). However, in the limit $\kappa M \ll V$ we get $n^* \sim \kappa \rho_0 - 1$, such that the organelle size is independent of cell size and the number of organelles assembled (Fig. 7e,f).

### X. EFFECT OF CELLULAR GROWTH ON ORGANELLE SIZE CONTROL

Here we briefly discuss the effect of cellular growth on the size control of organelles. We assume that a cell starts growing with an initial subunit abundance $N_0$ and volume $V_0$. For simplicity we assume a single rate of growth for both these variables -

$$\dot{V} = g$$
$$\dot{N} = g \tag{49}$$

where $g$ is the growth rate. Thus the timescale associated with cellular growth is $\tau_g = 1/g$. We implemented the growth of cell size and subunit abundance in a stochastic Gillespie simulation, and varied the timescale of organelle growth to study the effect cell growth on organelle size control affects. Specifically, we compare two distinct organelle growth mechanisms - (i) the limiting pool $\alpha + \beta = 0$, and the (ii) size-dependent growth model $\alpha + \beta = 1$. The timescales of these two growth mechanisms are $\tau_{LP} \sim \frac{1}{k^+}$ and $\tau_{\alpha\beta} \sim \frac{1}{k^+ + k^-}$, respectively.

In the limiting pool model (i), when organelle growth is much faster than cellular growth ($\tau_g \gg \tau_{LP}$) large anti-correlated size fluctuations persist and there is no control of individual organelle size (Fig. 8b). But when the organelle growth rate is comparable (or less than) to cellular growth ($\tau_g \sim \tau_{LP}$), transient control of organelle size is observed, with suppressed size fluctuations (Fig. 8a). In both cases, the individual mean size (over a time period $\sim \tau_{LP}$) increases as the total pool size increases in time. When there is competition between two non-identical structures, the faster growing structure takes up all the subunits an(d grows with cell size (Fig. 8b,inset). The size-dependent growth model (ii), however, ensures local temporal control of organelle size, with the mean size increasing with the growing cell size (Fig. 8c,d). Size control is ensured at both the limits $\tau_g \gg \tau_{\alpha\beta}$ and $\tau_g \sim \tau_{\alpha\beta}$.

---

[10] J. W. Raff, Trends in Cell Biology (2019).

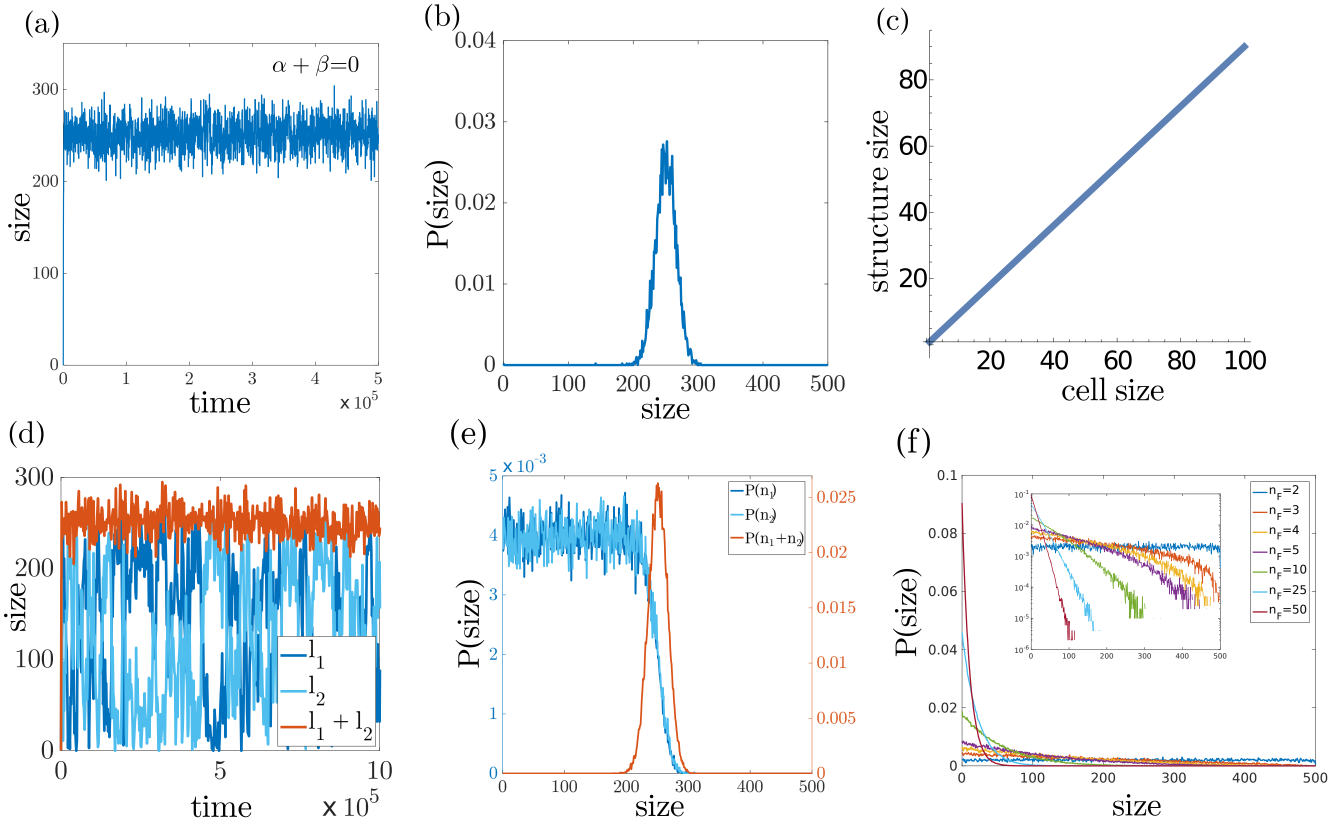


FIG. 1. Failure of the limiting pool in controlling the size of multiple structures. (a) For a single structure, the limiting pool model can provide robust size control. The structure reaches a steady state size after an initial period of fast growth. (b) The steady state size distribution shows a unimodal peaked distribution, characterising a well-defined mean size for the structure. (c) The limiting pool mechanism captures structure size scaling with cell size. (d-e) Limiting pool fails to control the individual size for two structures grown from a shared pool of subunits, giving rise to large anti-correlated fluctuations (d). The total size of the structures is a well controlled quantity, with temporal stability (d) and unimodal peaked distribution (e). The individual size distributions are almost uniform in a range of 0 to $N-\kappa^{-1}V$ (e). (f) For many structures, the individual size distributions converge to an exponential distribution - i.e., the standard deviation of size fluctuations are as large as the mean size, which is indicative of poor size control.

TABLE I. Parameter values for flagellar growth

| | | |
|---|---|---|
| tubulin concentration$(\rho_0) = 5\,\mu M$ | $\alpha = 1$ | $\beta = 0$ |
| tubulin size$(\delta L) = 10\,nm$ | $k^+ = 120\,\mu m^3\,min^{-1}$ | $k^- = 100\,min^{-1}$ |

TABLE II. Parameter values for *Long-zero* simulation

| | | | |
|---|---|---|---|
| tubulin concentration$(\rho_0) = 5\,\mu M$ | $t_0 = 1000\,min$ | $\alpha = 1$ | $\beta = 0$ |
| tubulin size$(\delta L) = 10\,nm$ | $k^+ = 120\,\mu m^3\,min^{-1}$ | $k^- = 100\,min^{-1}$ | $r_p = 0.0016\,min^{-1}$ |

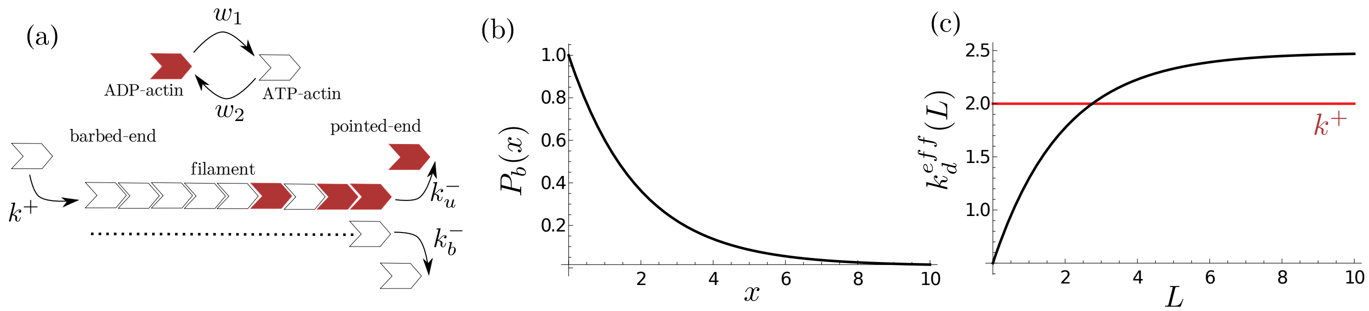


FIG. 2. Length dependent disassembly rate arises when monomers switch between different states with distinct disassembly rates. (a) Schematic of filament growth with monomer state switching. (b) At steady-state the probability of ATP-bound monomer $P_b(x)$ decreases towards the pointed end with a length scale $\lambda^{-1}$. (c) The effective length dependent disassembly rate $k_d^{eff}(L)$ can give rise to finite steady state length $\bar{L}$ as at this length $k^+ = k_d^{eff}(\bar{L})$. But it is important to notice that the maximum value for $k_d^{eff}$ is $k_{max} = k_u^- + A(k_b^- - k_u^-)$ and when $k^+ > k_{max}$ this effective length dependence cannot rescue the failure of size control and leads to uncontrolled growth of the filament. The parameter values used in the above results are: $k^+ = 2$, $k_b^- = 0.5$, $k_u^- = 2.5$, $w_1 = 0.01$ and $w_2 = 0.02$.

TABLE III. Parameter values for centrosome growth by uniform assembly and disassembly

| | | |
|---|---|---|
| subunit concentration$(\rho_0) = 1.67\,\mu M$ | $\alpha = -1$ | $\beta = 1$ |
| subunit size$(\delta V) = 5.8 \times 10^{-7}\,\mu m^3$ | $k^+ = 1.0\,\mu m^3\, min^{-1}$ | $k^- = 10^{-1}\, min^{-1}$ |

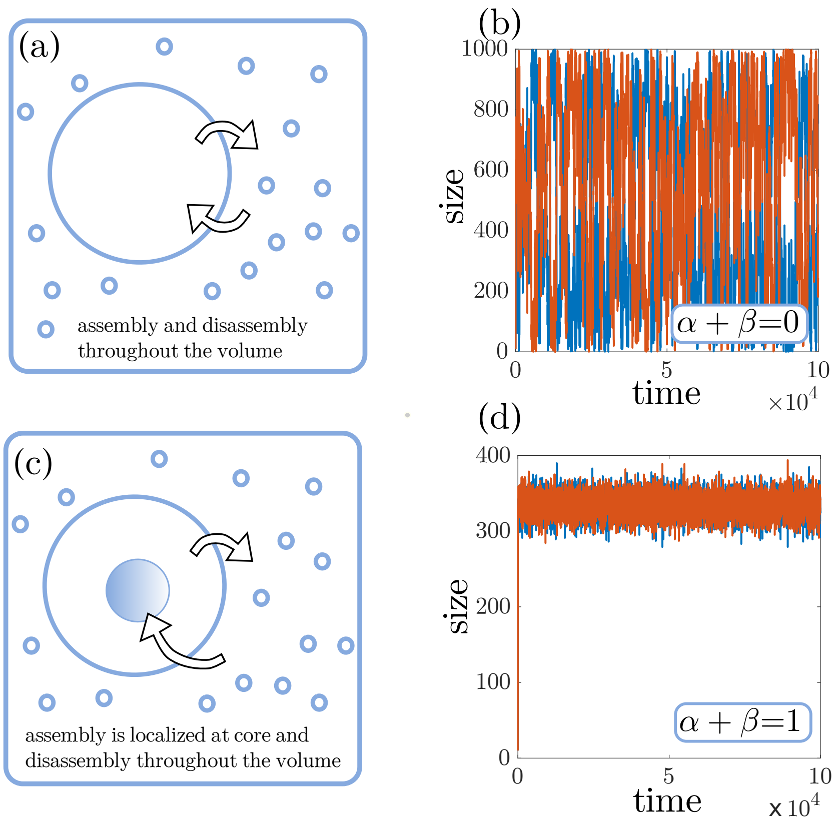


FIG. 3. Size control during centrosome growth. (a) Schematic of a sphere growing from a pool of subunits. The subunit assembly and disassembly can happen throughout the volume (porous structure) giving rise to a case where assembly rate and disassembly rate scale with volume of the structure -i.e., $\sim n$ where $n$ is the structure size in subunits. (b) A simple limiting pool cannot provide size control for multiple such spheres growing from a shared pool of subunits. (c) Schematic of a sphere growing from a fixed core. The assembly only occurs at the surface of the core by "chemical activity" of the core rendering a constant assembly rate independent of $n$. The disassembly can happen throughout the volume. (d) This localized reaction gives rise to effective size dependent negative feedback in growth and we obtain size control of individual spheres when two of them are growing from a shared pool.

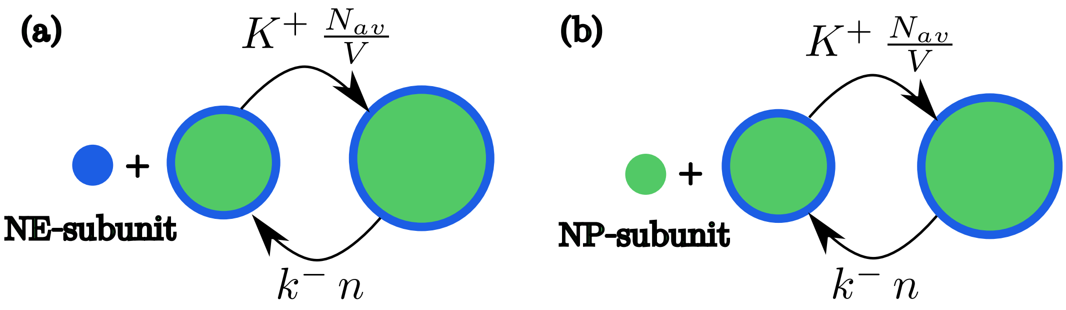


FIG. 4. Mechanisms of nuclear growth by nuclear envelope (NE) assembly and nucleoplasm (NP) assembly. (a) Nuclear envelope grows by incorporation of NE-subunits, and NP volume grows accordingly to accommodate the increase in nuclear surface area. (b) Nucleoplasm grows in volume by incorporation of NP-subunits, and the NE surface area increases accordingly to encapsulate the increase in NP volume.

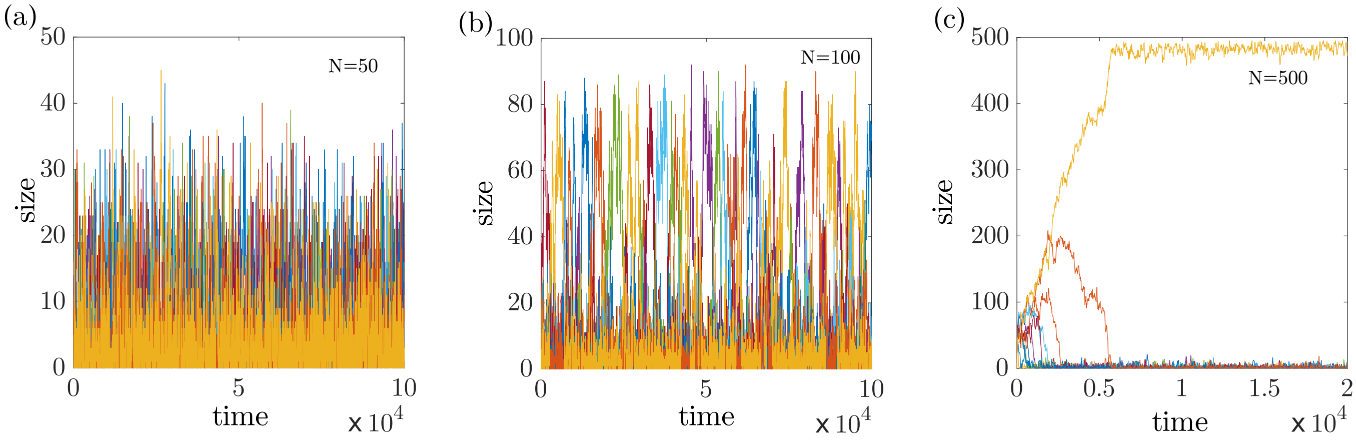


FIG. 5. Autocatalytic growth gives rise to bistability. (a-c) We study growth of multiple structures from a common pool of subunits, with size-dependent positive feedback $\alpha + \beta = -0.2$. This makes the growth process autocatalytic. We see a transition in the size dynamics as we increase the overall density of subunits (by changing total amount $N$ while keeping volume $V$ fixed). (a) The structures hardly grow at low subunit density. (b) At intermediate subunit density, bistability in size distribution emerges. The bigger structure captures most of the subunits, but suddenly starts declining in size due to stochastic fluctuations, when the other structures grow to be bigger. This creates a "flickering" growth pattern of multiple structures. (c) At a higher subunit density, we observe an initial growth of multiple structures. At a later time only a single structure forms while the others "die-out" in the competition. This mechanism can be used to make sure only a single structure gets built inside the cell which is important in various cases of polarity establishment and spontaneous symmetry breaking.

TABLE IV. Parameter values for centrosome growth by localised assembly and distributed disassembly

| | | |
|---|---|---|
| subunit concentration$(\rho_0) = 1.67\,\mu M$ | $\alpha = 0$ | $\beta = 1$ |
| subunit size$(\delta V) = 5.8 \times 10^{-7}\,\mu m^3$ | $k^+ = 10^3\,\mu m^3\,min^{-1}$ | $k^- = 10^{-3}\,min^{-1}$ |

TABLE V. Parameter values for centrosome growth: Two component model

| | | |
|---|---|---|
| $n_{a,b}$concentration$(\rho_0^{a,b}) = 1.67\,\mu M$ | subunit size$(\delta V) = 5.8 \times 10^{-7}\,\mu m^3$ | $\alpha = 0,\ \beta = 1$ |
| $k_a^+ = 12\,\mu m^3\,min^{-1}$ | $k_b^+ = 10\,\mu m^3\,min^{-1}$ | $k_{a,b}^- = 10^{-3}\,min^{-1}$ |

TABLE VI. Parameter values for nucleus growth by nuclear envelope assembly

| | | |
|---|---|---|
| NE subunit concentration$(\rho_0) = 8.0\,\mu m^{-3}$ | $\alpha = 0$ | $\beta = 1$ |
| NE subunit size$(\delta A) = 0.215\,\mu m^2$ | $k^+ = 5 \times 10^{-3}\,\mu m^3\,min^{-1}$ | $k^- = 10^{-1}\,min^{-1}$ |

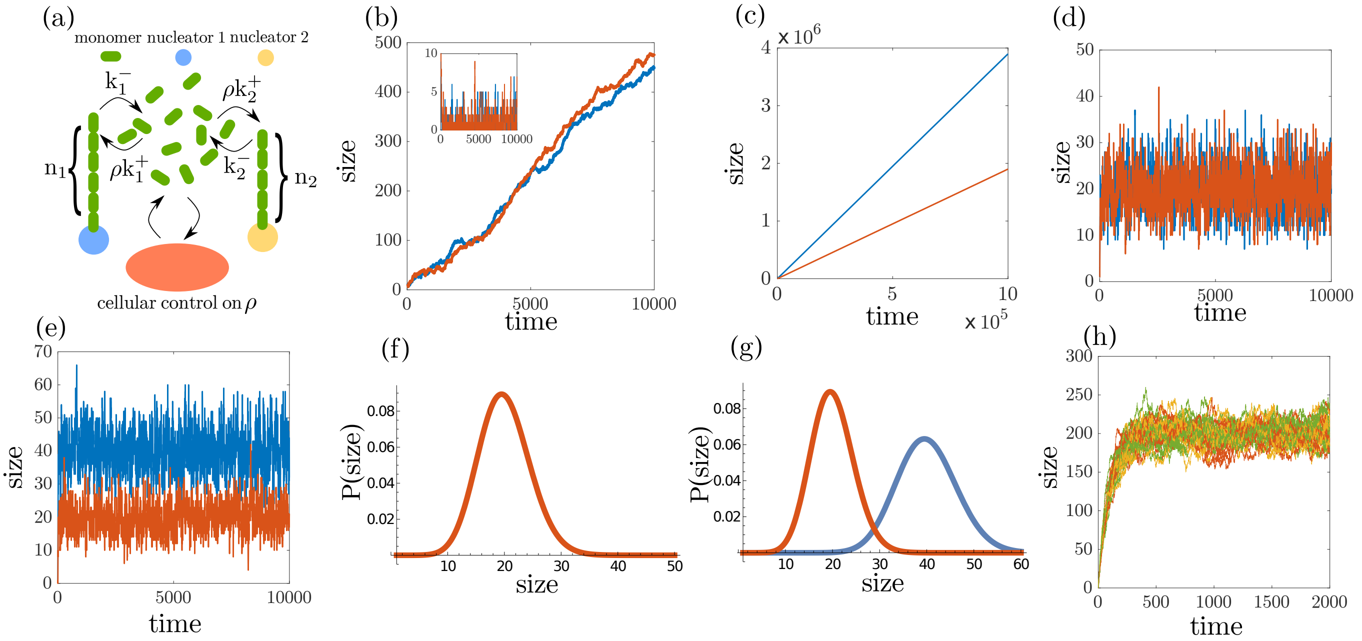


FIG. 6. Growth of structures under constant subunit concentration. (a) Schematic representation of the growth of two structures, where the cell maintains a fixed cytoplasmic density of subunits. (b-c) The structures grow in an unbounded manner when the concentration is higher than the critical concentration $\rho_c = \frac{k^-}{k^+}$. For $\rho < \rho_c$ the structures do not grow (b, inset). Here $\kappa_1 = \kappa_2 = 10$ and $\rho = 0.15$ in (b), and $\kappa_1 = 2\kappa_2 = 200$ and $\rho = 0.2$ in (c). (d-e) Size-dependent negative feedback enables robust size control of two structures with (e) or without (d) competition. (f-g) The steady-state size distributions for the two structures in (d) and (e), computed analytically by solving the master equation. (h) The constant subunit concentration model does not lead to any scaling between the structure size and the system size, when multiple structures growing from a shared subunit pool. The plot shows time series for the size of 10 (blue), 20 (yellow) and 40 (red) structures grown from a pool of concentration $\rho = 2$, with $\kappa = 100$.

TABLE VII. Parameter values for nucleus growth by nucleoplasm assembly

| | | |
|---|---|---|
| NP subunit concentration$(\rho_0) = 0.75\,\mu m^{-3}$ | $\alpha = 0$ | $\beta = 1$ |
| NP subunit size$(\delta V) = 0.1\,\mu m^3$ | $k^+ = 2.0\,\mu m^3\,min^{-1}$ | $k^- = 10^{-3}\,min^{-1}$ |

TABLE VIII. Parameter values for microtubule aster dynamics in nucleus growth

| | | |
|---|---|---|
| MT subunit concentration$(\rho_0) \simeq 0.67\,\mu M$ | $\alpha = 0$ | $\beta = 1$ |
| MT subunit size$(\delta L) = 5\,nm$ | $k_m^+ = 2.0\,\mu m^3\,min^{-1}$ | $k_m^- = 10^{-2}\,min^{-1}$ |

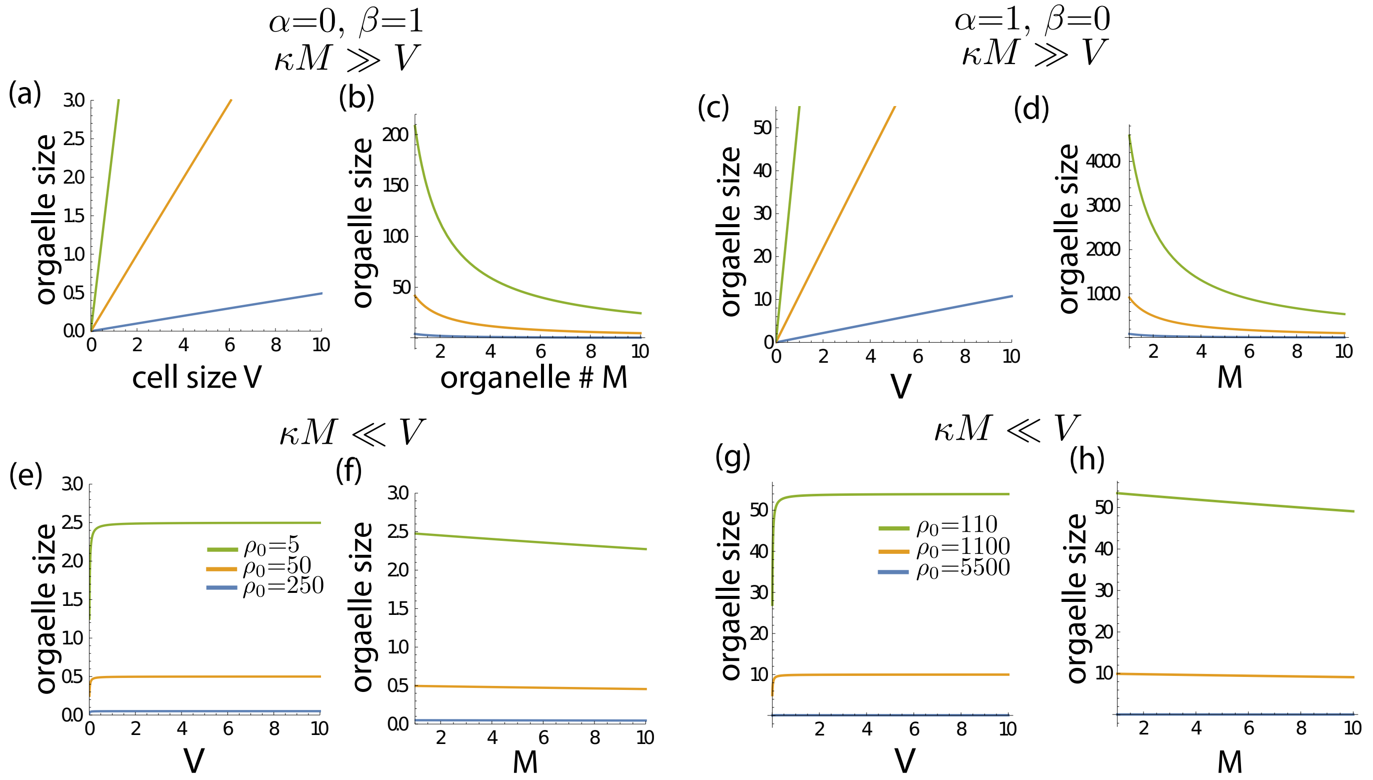


FIG. 7. Organelle-to-cell size scaling regimes. The size-dependent growth model (with $\alpha = 0$, $\beta = 1$ and $\alpha = 1$, $\beta = 0$) exhibits a regime where organelle size does not scale with cell/system size ($V$) or the number of assembled organelles, $M$. Organelle growth rate can be tuned to achieve scaling of organelle size with cell size. (a-d) shows the organelle-to-cell size scaling regimes for $\alpha = 0$, $\beta = 1$ and (e-h) shows the same regimes for $\alpha = 1$, $\beta = 0$.

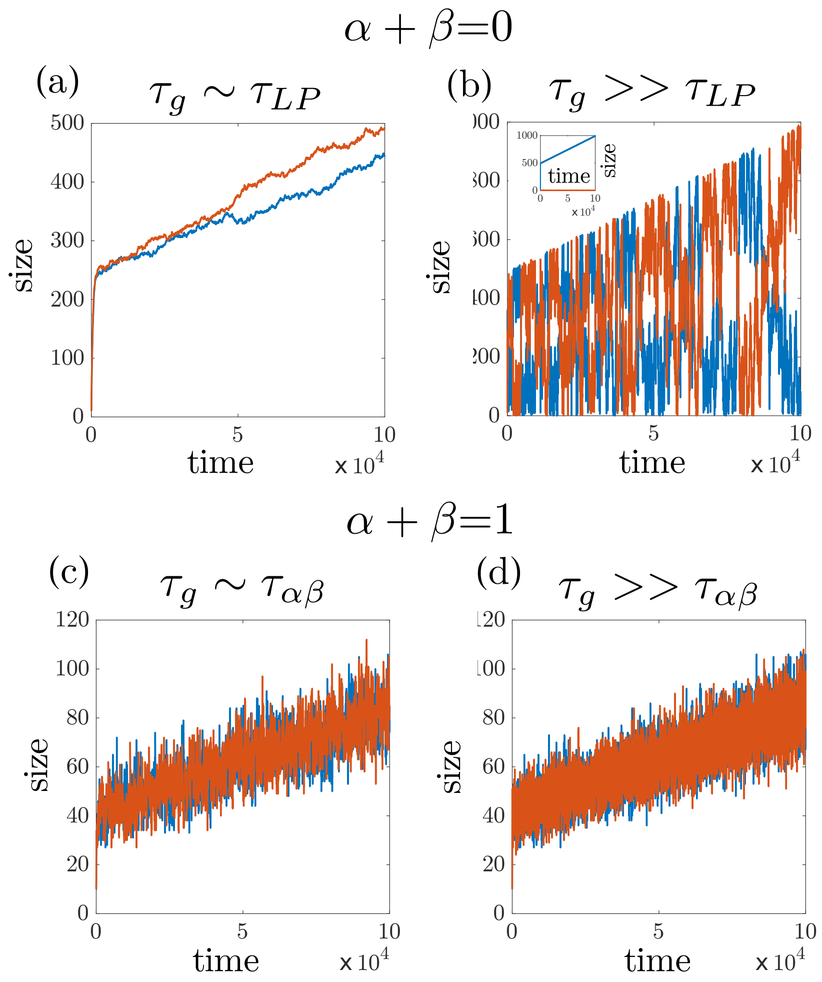


FIG. 8. Effect of cellular growth on organelle size control. (a,b) In the limiting pool model ($\alpha + \beta = 0$), fast organelle growth shows characteristic large fluctuations and loss of size control. In the slow growth regime for organelles, transient size control is observed without significant size fluctuations. (c,d) The size-dependent growth model ($\alpha + \beta = 1$) provides local temporal control of organelle size, with the organelle size growing at the same rate as cellular growth.
